# Requirement of *Irf6* and *Esrp1/2* in frontonasal and palatal epithelium to regulate craniofacial and palate morphogenesis in mouse and zebrafish

**DOI:** 10.1101/2020.06.14.149773

**Authors:** Shannon H. Carroll, Claudio Macias Trevino, Edward B-H Li, Kenta Kawasaki, Nora Alhazmi, Shawn Hallett, Justin Cotney, Russ P. Carstens, Eric C. Liao

## Abstract

Orofacial clefts are among the most common human congenital malformations. *Irf6* and *Esrp1* are two key genes important for palate development, conserved across vertebrates. In the zebrafish, we found that irf6 regulates the expression of *esrp1*. Using RNAscope, we detailed overlapping *Irf6* and *Esrp1/2* gene expression in the mouse frontonasal prominence ectoderm, lambda joint periderm, palate and lip epithelium. In the zebrafish, *irf6* and *esrp1/2* share expression in the pre-gastrulation periderm and the embryonic frontonasal ectoderm, oral epithelium ventral to the anterior neurocranium (ANC), and the developing stomodeum. Genetic disruption of *irf6* and *esrp1/2* in the zebrafish resulted in cleft of the ANC. In the *esrp1/2* zebrafish mutant, cleft of the mouth opening formed and appeared to tether into the ANC cleft. Lineage tracing of the anterior cranial neural crest cells revealed that cleft of the ANC resulted not from migration defect, but from impaired chondrogenesis. Molecular analysis of the aberrant cells localized within the ANC cleft revealed that this cell population espresses *sox10, col1a1* and *irf6* and is adjacent to cells expressing epithelial *krt4*. Detailed morphogenetic analysis of mouse *Irf6* mutant revealed mesenchymal defects not observed in the *Esrp1/2* mutant. Analysis of breeding compound *Irf6;Esrp1;Esrp2* mutant suggests that these genes interact where the triple mutant is not observed. Taken together, these studies highlight the complementary analysis of *Irf6* and *Esrp1/2* in mouse and zebrafish models and captured an unique aberrant embryonic cell population that contributes to cleft pathogenesis. Future work characterizing this unqiue *sox10+, col1a1+, irf6*+ cell population will yield additional insight into cleft pathogenesis.

## INTRODUCTION

The development of vertebrate craniofacial structures requires coordinated cellular induction, migration, proliferation, and differentiation, which allow for the positioning of adjacent epithelial-lined facial processes that ultimately merge(Abramyan and Richman, 2015; Cordero et al., 2011; Creuzet et al., 2005; Dougherty et al., 2012; Helms et al., 2005; Jiang et al., 2006; Knight and Schilling, 2006; Mork and Crump, 2015; Reid et al., 2011). Morphogenesis of facial structures such as the midface, lip, and palate requires convergence of the medial and lateral nasal prominences and the fusion of the secondary palatal shelves at the midline (Abramyan and Richman, 2015; Jiang et al., 2006; Losa et al., 2018). Failure of these processes to fuse results in orofacial clefts (OFCs) of the lip, primary palate, or secondary palate (Gritli-Linde, 2008). Orofacial clefts are among the most common congenital structural anomalies (Goodwin et al., 2015; Juriloff and Harris, 2008; Yuan et al., 2011). From genome-wide association studies over a decade ago to more recent whole-genome sequencing projects of orofacial cleft cohorts, human geneticists have identified several cleft-associated genetic loci, where the transcription factor *IRF6* continues to be one of the most commonly associated genes (Marazita, 2012; Zucchero et al., 2004). *IRF6* disruption is causal for syndromic cleft in Van der Woude and Popliteal Pterygium syndromes, and associated with non-syndromic orofacial clefts (Beaty et al., 2016; Kondo et al., 2002; Leslie et al., 2013; Zucchero et al., 2004).

Several IRF6 transcriptional targets such as *GRHL3*, *WDR65*, *OVOL1, KLF4* have been identified, which are also important for palate development and implicated in human cleft pathogenesis (de la Garza et al., 2013; Kousa and Schutte, 2016; Liu et al., 2016; Rorick et al., 2011). These studies support the premise that investigation of *Irf6* and its transcriptional network will identify key genes that regulate palate development. Multiple mouse models have been generated to investigate *Irf6* function, including a total *Irf6* knockout as well as amino acid substitution of key functional residue *Irf6*^*R84C*^ in the DNA-binding domain (Ingraham et al., 2006; Richardson et al., 2006). These *Irf6* mutant mice exhibited disrupted epithelial terminal differentiation and lack of a functional periderm, leading to pathological adhesions of epithelial tissues (Ferretti et al., 2011; Ingraham et al., 2006; Iwata et al., 2013; Richardson et al., 2006). The epithelial differentiation and adhesion defects are thought to prevent elevation of the palatal shelves, and ultimately these mice have a cleft in the secondary palate. Additionally, the midface of these mice were hypoplastic, which was attributed to a dysfunctional epithelium (Ingraham et al., 2006; Richardson et al., 2006).

*Epithelial splicing regulatory factors 1* and *2* (*Esrp1*, *Esrp2*) are also important in embryonic epithelial differentiation and palate development (Bebee et al., 2015; Lee et al., 2018; Lee et al., 2020). *Esrp2* and its homolog *Esrp1* are regulators of RNA splicing that are specifically expressed in the epithelium (Warzecha et al., 2009). *Esrp1/2* knockout mice exhibit bilateral cleft of the lip and primary palate, as well as a secondary palate cleft (Bebee et al., 2015). *Esrp1/2* are unusual among regulators of RNA-splicing in that they are tissue-restricted and exhibit dynamic expression during embryogenesis (Bebee et al., 2015; Burguera et al., 2017). The developmental importance of *Esrp1/2* is underscored by their conservation across species, from ascidians to zebrafish, *Xenopus*, mouse, and humans (Burguera et al., 2017). Gene variant in *ESRP2* was also recently reported in human orofacial cleft cohorts (Cox et al., 2018).

The mouse has been an important experimental model to study craniofacial and palate development (Gritli-Linde, 2008). Secondary palate development in the mouse is similar to humans, with the analogous stages of vertical outgrowth, elevation, horizontal growth, and fusion (Gritli-Linde, 2008; Juriloff and Harris, 2008). Many genes associated with cleft lip and palate (CL/P) in humans when disrupted in the mouse result in cleft of the secondary palate, but the primary palate and lip appear unaffected (Gritli-Linde, 2008; Van Otterloo et al., 2016). So while the mouse model can be useful to study the secondary palate, the use of mouse models to study cleft of the lip and primary palate has been less effective as there are remarkably few mouse models where development of the lip and primary palate are perturbed (Gritli-Linde, 2008). Meanwhile, clinically CL/P is more common than isolated cleft of the palate only (CPO), and human genetic studies have suggested that the genetics underpinning CL/P and CPO are distinct (Gritli-Linde, 2008; Juriloff and Harris, 2008) The developmental processes of outgrowth of the facial prominences followed by convergence and fusion are thought to be conserved across mammals (Abramyan and Richman, 2015). Therefore it is hypothesized that differences in mouse versus human phenotypic presentation are due to spatiotemporal differences in craniofacial development (Gritli-Linde, 2008). In this context, the phenotype of bilateral clefts affecting the lip, primary and secondary palate in the *Esrp1/2* mutant mouse is unique among mouse models and is a valuable tool to study lip and palate morphogenesis.

Zebrafish has been favored as an animal model by embryologists to study craniofacial development due to its accessibility and transparency (Kimmel, 1989; Lieschke and Currie, 2007; Schilling and Le Pabic, 2009). Although a secondary palate, which partially or entirely separates the oral and nasal cavities, is reserved to select amniotes, the primary palate is appreciably conserved across all vertebrates (Abramyan and Richman, 2015). The primary palate establishes the intact upper jaw (Abramyan and Richman, 2015), which in the larval zebrafish consists of the ethmoid plate, also known as the anterior neurocranium (ANC). In all vertebrates, the most anterior CNCCs that migrate rostral then turn caudal and ventral to the eye contribute to the median frontonasal prominence, and a second CNCC stream that migrates or inferior to the eye and into the first pharyngeal arch to generate the paired maxillary prominences (Dougherty et al., 2012; Kimmel, 1989; Swartz et al., 2011; Wada et al., 2005). The ANC of the zebrafish is formed from the convergence of the median element that is derived from the frontonasal prominence, and paired lateral elements that are derived from the maxillary prominences (Duncan et al., 2017; Mork and Crump, 2015; Swartz et al., 2011). Zebrafish homologs of human genes associated with orofacial clefts will disrupt morphology of the ANC, as have been observed fro a number of genes such as *capzb*, *pitx2*, *pdgfra*, *smad5*, *tgfb2*, *fgf10a* and wnt9a (Duncan et al., 2017; Mork and Crump, 2015; Van Otterloo et al., 2016).

Here, we carried out detailed gene expression analysis of *Irf6*, *Esrp1* and *Esrp2* in mouse and zebrafish in order to understand the comparative morphogenesis of facial structures and periderm between these common vertebrate genetic models. We analyzed and compared the *Irf6* and *Esrp1/2* mutant phenotypes to elucidate the comparative morphologies and genetic epistasis between these genes. Further, we generated zebrafish *irf6* and *esrp1/2* zebrafish mutants and examined their requirement in morphology of the stomodeum opening and ANC. Interestingly, we identified an aberrant cell population with epithelial and mesenchymal molecular signatures that localized to the region of the ANC cleft. This work highlights the relative strengths of the mouse and zebrafish models for investigating the morphogenetic mechanisms of orofacial clefts and contributes new insights into the function of *Irf6* and *Esrp1/2* during palatogenesis.

## RESULTS

### *irf6* null zebrafish embryos have decreased expression of *esrp1*

We previously reported generation of a functionally null *irf6* zebrafish allele (Li et al., 2017). Using CRISPR/Cas9, an 8 bp deletion in exon 6 of the *irf6* coding region resulting in a frameshift and premature stop codon, leading to the ablation of *irf6* function. It was observed that embryos lacking maternally expressed *irf6* exhibited epiboly arrest and periderm rupture at 4-5 hpf (Li et al., 2017). Utilizing this maternal *irf6*-null model, we aimed to identify genes that were differentially downregulated in *irf6*-null versus wild type embryos. We performed RNA-seq on wild type and maternal/zygotic *irf6*-null (mz-*irf6*^−8bp/−8bp^) embryos at 4.5 hpf, just before embryo rupture at the onset of gastrulation, as *irf6* is required for periderm integrity. Analysis of the RNA-seq identified differentially-expressed genes (DEGs) in wild type relative to mutant (Fig. 1A-C). The RNA-seq results revealed significant downregulation of genes previously known to be downregulated with disruptions in *Irf6* function (Fig.1B,C). Disruption of *irf6* via injecting dominant-negative *irf6* mRNA led to downregulation of many periderm-enriched genes (including *grhl1*, *krt5*, *krt18*, *tfap2a* and *klf2b*) and genes for adhesion molecules (including claudins and cadherins) (de la Garza et al., 2013). Here we found a similar expression profile in the mz-*irf6*^−8bp/−8bp^ embryos relative to wild type (Fig. 1B,C).

**Figure 1.**
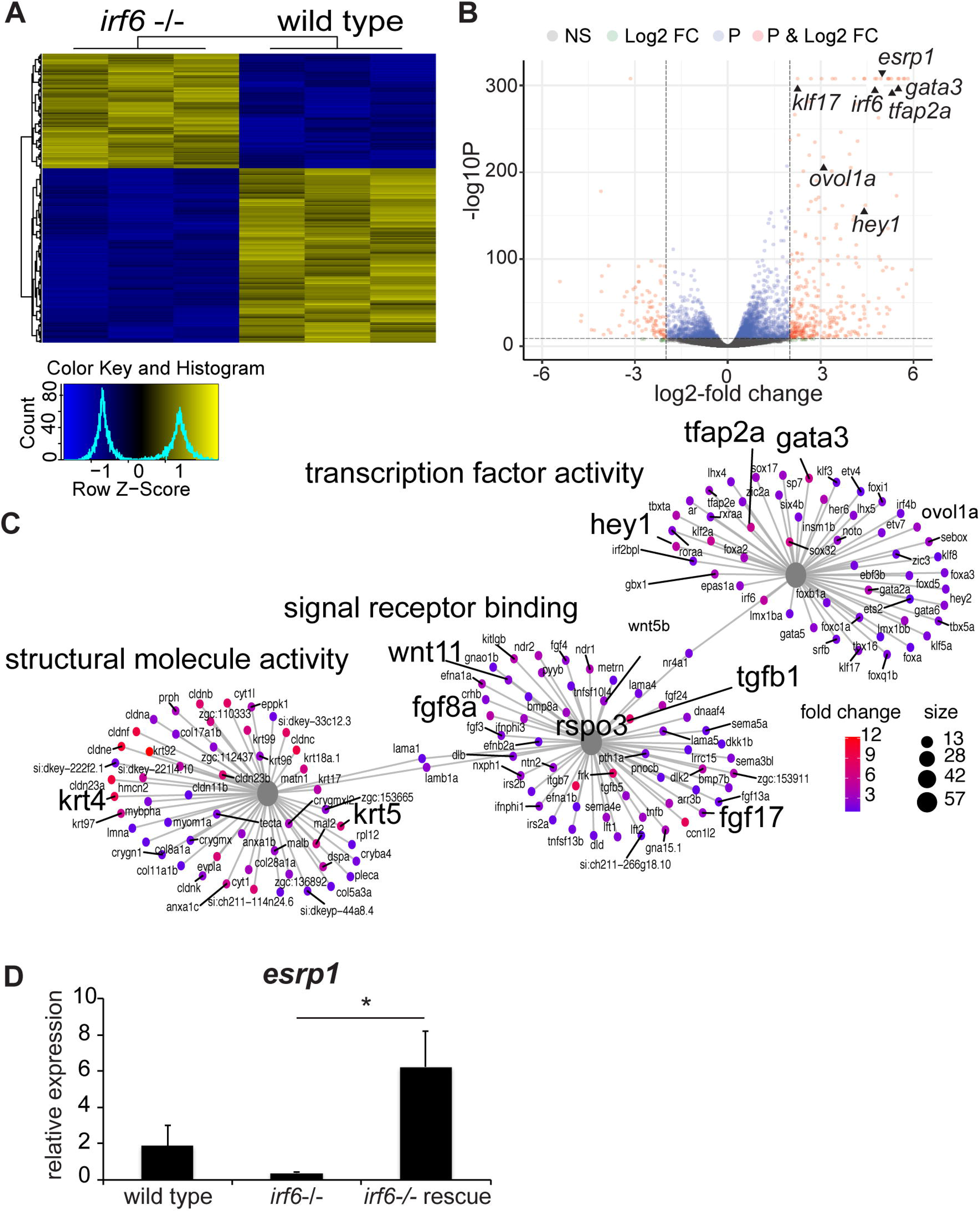
*esrp1* expression is downregulated in *irf6* null zebrafish embryos. (A) Hierarchical clustering of top differentially expressed genes (DEGs) defined by RNA-sequencing performed on WT vs. mz-*irf6*^*−8bp/−8bp*^ (*irf6−/−*) zebrafish embryos at 4-5hpf. Top DEGs were identified by selecting genes with an adjusted p-value (Benjamini-Hochberg) <0.01 and absolute log2-fold change > 2. Data shown for 3 biological replicates. Color scale at top left represents relative levels of expression with yellow showing higher expression levels and blue showing lower expression. (B) Volcano plot from the RNA-seq dataset showing the distribution of DEGs based on p-values and log2-fold change. Previously published *irf6*-regulated genes are expressed at significantly higher level in wild type relative to mz-*irf6*−/−, including *grhl3*, *klf17,* and *wnt11*. The newly identified cleft-associated gene *esrp1* is also expressed significantly higher in wild type relative to *irf6*−/−. Horizontal and vertical lines represent the p-value cutoff of 0.01 and the log2-fold change cutoff of 2, respectively. (C) Gene ontology (GO) gene-concept network analysis of RNA-seq data showing that *irf6*−/− embryos have perturbations in processes such as transcription factor activity, signal receptor binding, and structural molecule activity. Note that many of these genes, such as *wnt11*, *fgf8*, *tgfb1*, *krt4*, and *krt5*, are implicated in ectoderm development and cell specification. Grey nodes show GO terms, colored nodes show individual genes from the RNA-seq dataset, and edges connect genes to one or more associated GO terms. Colored nodes show relative enrichment (measured by fold-change) of genes in wild type samples relative to *irf6−/−* embryos. Maps were generated using the enrichplot package in R. (D) qPCR gene expression analysis for *esrp1*, showing approximately 6-fold *esrp1* downregulation in mz-*irf6*^−8bp/−8bp^ embryos compared to wild type at 4 hpf, and rescued *esrp1* gene over-expression in mz-*irf6*^−8bp/−8bp^ embryos injected with wild type zebrafish *irf6* mRNA. n = 4. *p<0.05.

When compared to previously published *IRF6* siRNA human keratinocyte DEG expression data (Botti et al., 2011), there were major overlaps of genes in molecular pathways responsible for epithelial regulation, including *gata3*, *krt18* and *cldn4* (Fig.1 B,C and Fig. S1).

In addition to the epithelial differentiation and maturation genes, many key developmental signaling pathways including Fgf (*fgf8a*, *fgf17* and *fgf24*) and Wnt (*wnt11*, *dact2*, *rspo3*, *frzb*, *fzd5*) pathways were also heavily represented in our dataset as genes downregulated due to *irf6* ablation (Fig. 1B,C). Further, a number of genes associated with human orofacial clefts (OFCs) are also downregulated in the *irf6* null embryos, including *hey1*, *gata3*, *wnt11*, and *fgf8* (Fig. 1B,C).

Interestingly, one of the most downregulated genes in the mz-*irf6*^−8bp/−8bp^ embryos was *esrp1*. *Epithelial splicing regulatory protein 1* (*esrp1)* and its paralog *esrp2* are epithelial restricted RNA splicing regulators. *ESRP2* genetic variants in humans are associated with OFCs (Cox et al., 2018), and *Esrp1* and *Esrp1/2* knockout mice display a bilateral cleft of the lip, primary and secondary palate (Bebee et al., 2015). To confirm the RNA-seq results, we performed qPCR on mz-*irf6*^−8bp/−8bp^ and wild type embryos at 4-5 hpf. Relative to wild type, mz-*irf6*^−8bp/−8bp^ embryos had approximately 5-fold downregulation of *esrp1* expression. Additionally, injection of mz-*irf6*^−8bp/−8bp^ embryos with *irf6* mRNA at the 1-cell stage rescued *esrp1* expression, resulting in increase that was approximately 3-fold higher than wild type (Fig. 1D). Therefore *esrp1* gene expression is dependent on *irf6*, either through direct regulation or the requirement of a normal periderm.

### *irf6*, *esrp1* and *esrp2* are co-expressed in the oral epithelium of zebrafish during craniofacial development

Previous mouse studies have described *Irf6*, *Esrp1,* and *Esrp2* gene expression in epithelial cells during palate development (Bebee et al., 2015; Knight et al., 2006). To determine the gene expression of *irf6* and *esrp1* and *esrp2* in the zebrafish during epithelial and craniofacial development, we performed whole-mount *in situ* hybridization (WISH) as well as RNAscope *in situ* hybridization (ISH) at 8-cell stage, shield stage, 48 and 72 hpf. WISH of zebrafish embryos demonstrated maternal deposition of *irf6*, *esrp1* and *esrp2* mRNA detectable at 8-cell stage (Fig. 2A). The maternal transcriopt of these genes are also detected in the periderm of the gastrulating embryo, although expression of *esrp2* appears lower than *irf6* and *esrp1* (Fig.2A). During craniofacial development WISH demonstrated specific expression of *irf6*, *esrp1,* and *esrp2* lining the embryonic oral epithelium, and circumscribing surface epithelium concentrated around the developing stomodeum (Fig. 2B,C). Gene expression domains of *irf6*, *esrp1,* and *esrp2* overlap significantly, consistent with previous reports (Burguera et al., 2017).

**Figure 2.**
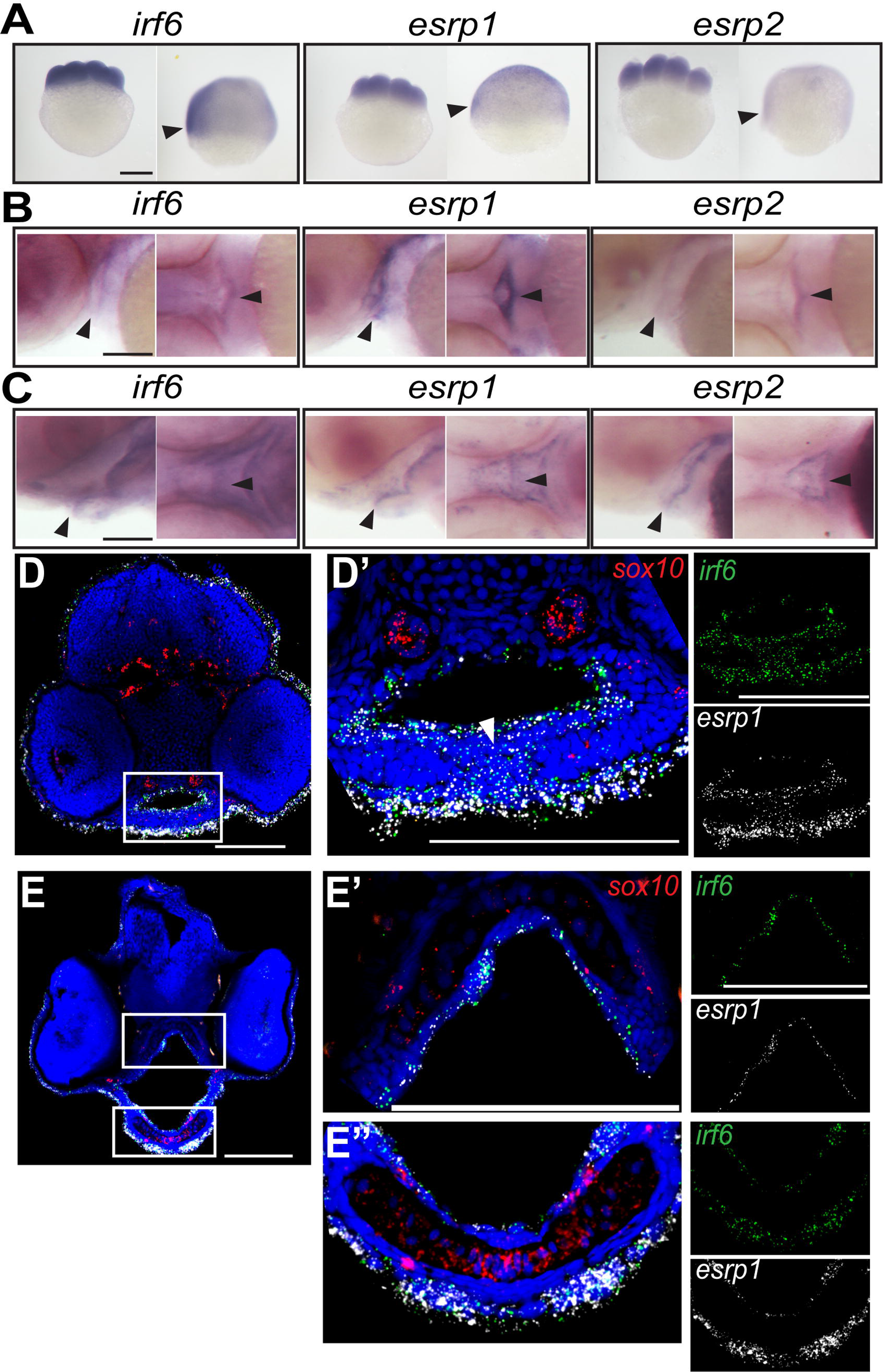
*irf6, esrp1* and *esrp2* are co-expressed in the oral epithelium of zebrafish embryos. (A-C) Whole mount *in situ* hybridization showing *irf6*, *esrp1* and *esrp2* maternal deposited transcripts are detected at the 8-cell and shield stage (A; arrow depicts periderm) circumscribes the developing stomodeum and lines the oral epithelium of zebrafish embryos at 48 (B) and 72 (C) hpf (depicted by arrow). All whole-mount embryos are oriented with anterior toward left of page, dorsal toward top of page. Coronal sections of 48 (D) and 72 (E) hpf embryos analyzed by RNAscope *in situ* hybridization, (dorsal toward top of page) showing cellular RNA co-expression of *irf6* (green) and *esrp1* (white) in surface and oral epithelial cells. *sox10* (red) staining depicts cartilage elements of the palate. White arrow depicts area of fusion between Meckel’s cartilage elements. Scale bars: 250μm (A) and 100μm (B-E).

To resolve the specific cell population of the embryonic epithelium that express *irf6*, *esrp1,* and *esrp2*, we performed RNAscope *in-situ* hybridization of coronal cryosections taken through the developing mouth and palate at 48 and 72 hours post-fertilization (hpf). We find that *irf6* and *esrp1* are co-expressed within epithelial cells lining the oral cavity as well as the surface epithelium (Fig. 2D,E). No expression of these genes was detected within the cartilage elements, identified by *sox10* expression. Further, we detected *irf6* and *esrp1* transcripts within the same cells, importantly within epithelial cells separating adjacent cartilage elements prior to the fusion of paired Meckel’s cartilage elements derived from the mandibular facial prominence (Fig. 2G’).

### *Irf6*, *Esrp1,* and *Esrp2* are co-expressed in murine frontonasal and oral epithelium, during palate and lip development

*Irf6*, *Esrp1, and Esrp2* are each expressed within the embryonic oral epithelium surrounding the developing palatal shelves of mice (Bebee et al., 2015; Knight et al., 2006). Ablation of *Irf6* or *Esrp1/2* causes a cleft of the secondary palate, but the disruption of the lip and primary palate phenotypes differ between the *Irf6* and *Esrp1* and *Esrp2* mutants (Bebee et al., 2015; Ingraham et al., 2006; Richardson et al., 2006). To determine whether *Irf6, Esrp1* and *Esrp2* transcripts co-localize during mouse craniofacial development similarly to zebrafish, we performed whole-mount ISH for each gene starting at E10.5 as the frontonasal prominences and lambdoidal junction are taking shape at this time point. We found that *Irf6*, *Esrp1,* and *Esrp2* genes are expressed similarly in E10.5 embryos with high levels of expression in areas of craniofacial development (Fig. 3A). The mouse gene expression pattern was similar to that observed in zebrafish, with more concentrated expression to the developing head. Higher-resolution imaging with RNAscope ISH to determine cellular co-localization detected *Irf6*, *Esrp1,* and *Esrp2* transcripts in the periderm and the basal epithelium across all time points examined (Fig. 3B-F). *Irf6*, *Esrp1* and *Esrp2* are co-expressed in the surface ectoderm overlying the developing frontonasal prominences (Fig. 3B), a cell population with important signaling and inductive functions (Hu et al., 2003). Further, co-expression included cells at critical fusion points, specifically between the medial and lateral nasal prominences (Fig.3C) and the palatal shelves (Fig. 3E,F). Additionally, with RNAscope, we were able to resolve differences in the mRNA expression pattern of *Esrp1* and *Esrp2*, with *Esrp2* generally being more highly expressed in the apical epithelial layer. In contrast, *Esrp1* is evenly expressed throughout the apical and basal epithelial layer. Interestingly, in addition to *Irf6* expression in the epithelium, RNAscope detected *Irf6* mRNA expression in the mesenchyme, particularly at E10 and E15.5 (Fig. 3B,E). Expression of *Irf6* in this craniofacial mesenchyme has not been reported. However, expression was detected in CNCCs of the first and second pharyngeal arches at E9.0 (Fakhouri et al., 2017) and *Irf6* is expressed in cells of the developing tongue (Goudy et al., 2013). Further, we previously reported that zebrafish expressing the *Irf6*^*R84C*^ variant under a *sox10* promoter exhibit a partial cleft of the anterior neurocranium (Dougherty et al., 2013). Together, these results suggest a role of *Irf6* in craniofacial development beyond its role in epithelial cell differentiation and fusion.

**Figure 3.**
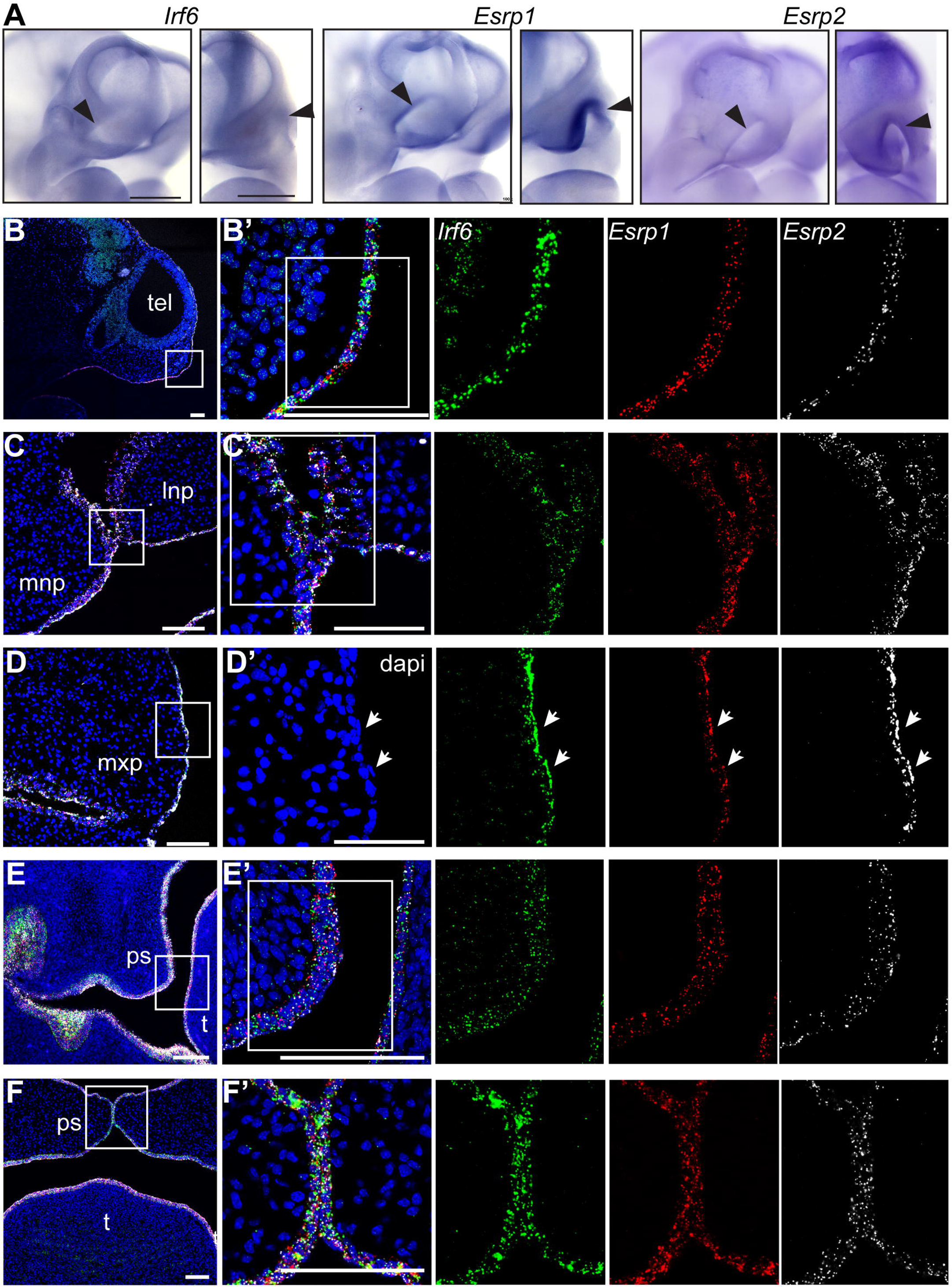
*Irf6*, *Esrp1* and *Esrp2* are co-expressed in the oral epithelium of mouse embryos. (A) Whole mount *in situ* hybridization of E10.5 embryos showing *Irf6*, *Esrp1* and *Esrp2* mRNA expression in the surface epithelium and concentrated within the ectoderm of the frontonasal prominences and first brachial arch. Oblique and frontal orientation. Scale bars: 500 μm. (B-F) Sections of (B) E10, (C,D) E11.5, (E) E13.5 and (F) E15 embryos analyzed by RNAscope *in situ* hybridization showing mRNA cellular co-expression of *Irf6* (green), *Esrp1* (red) and *Esrp2* (white) in the surface ectoderm (E10), lining the frontonasal and maxillary prominences, including expression in periderm (arrows) (E11.5) and lining the palatal shelves (E13.5, E15). (B) sagittal and (C-F) coronal sections. Scale bars: 100 μm.

### Disruption of *irf6* during neural crest cell migration results in cleft palate in zebrafish

Germline mutation of *irf6* results in early embryonic lethality due to periderm rupture during gastrulation, which precluded evaluation of palate morphogenesis that occurs later in development (Li et al., 2017; Sabel et al., 2009). To circumvent embryonic lethality, we employed an optogenetic gene activation system based on the light-sensitive protein EL222, that serves to induce the expression of genes downstream of the C120 promoter (Motta-Mena et al., 2014). To this end, a dominant-negative form of *irf6* consisting of a fusion protein of the *irf6* protein-binding domain and the engrailed repressor domain (*irf6*-ENR) was cloned downstream of the C120 promoter (C120-*irf6*-ENR; Fig. 4A) (Sabel et al., 2009). When co-injected with VP-16 mRNA, this light activated *irf6*-ENR construct enabled us to control the timing of *irf6* disruption by exposing the embryos to a 465nm light-source later in embryogenesis, thereby circumventing gastrulation lethality in the *irf6* mutants (Fig. 4A). Zebrafish embryos that were injected with the optogenetic system and continuously exposed to blue light from 10-96 hpf were able to survive, but developed with a slightly curved body axis and a dysmorphic ventral cartilage phenotype, which were not observed in control injected embyros or injected embryos that were raised in the dark (Fig. 4B). Further analysis of the cartilage in these embryos through Alcian blue staining and micro-dissections revealed a cleft in the ANC where a population of cells in the median portion of the ANC was absent (Fig.4C).

**Figure 4.**
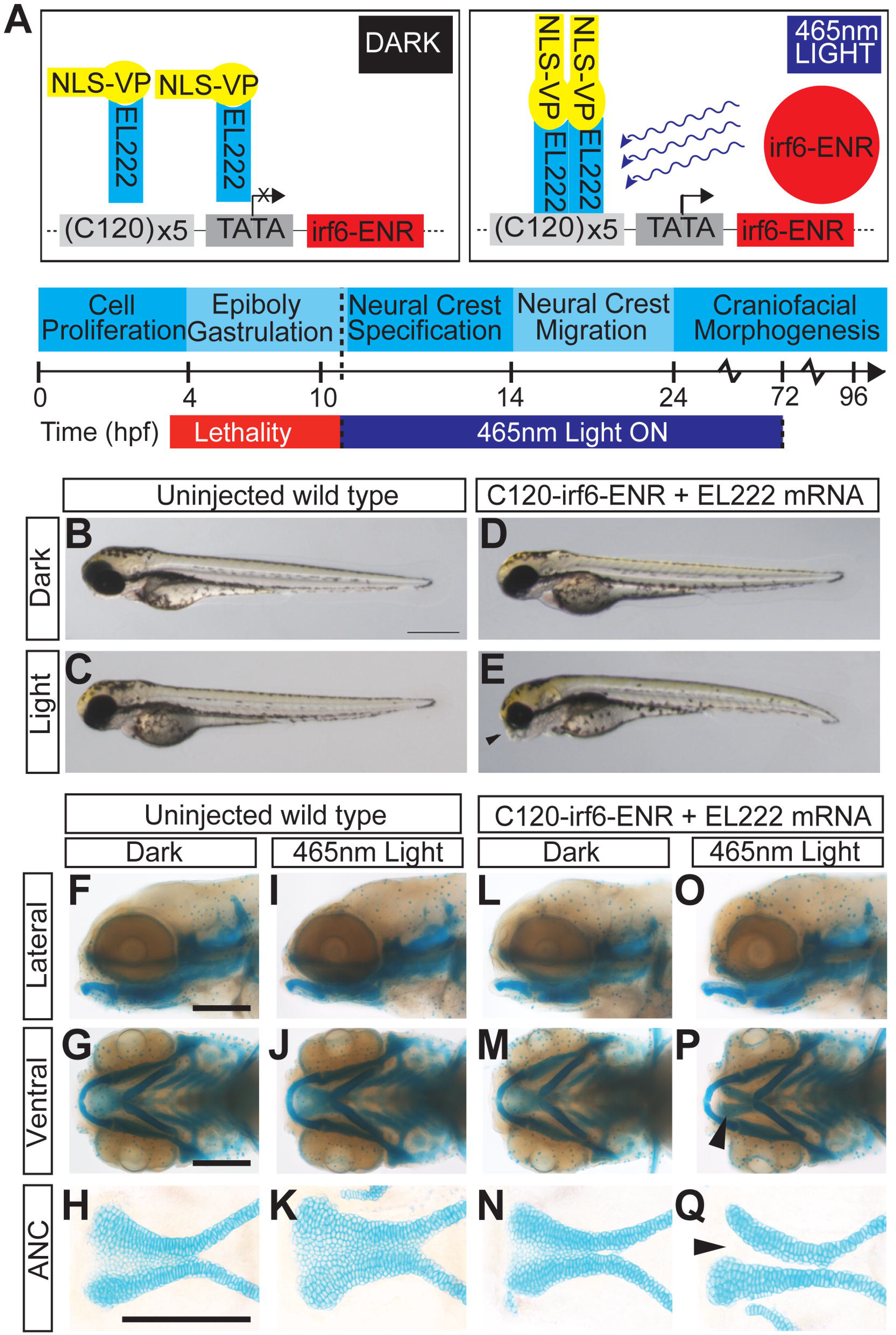
EL222 optogenetic disruption of *irf6* circumvents early embryonic lethality and causes a cleft palate phenotype. (A) Schematic of EL222 Optogenetic system. VP16-EL222 monomers are inactive under dark conditions. Upon stimulation by 465nm light, VP16-EL222 dimerizes, drives gene expression downstream of the C120 promoter, and induces the expression of a dominant negative form of irf6 (*irf6*-ENR). Embryos were exposed to blue light from 10-72 hpf to circumvent embryonic lethality in mz-*irf6*^−8bp/−8bp^ embryos. (B-E) Brightfield microscopy of 72 hpf zebrafish embryos injected with the optogenetic system and grown in the dark (D) or exposed to blue light from 10-72 hpf (E) compared to control injected embryos (B, C). Injected fish exposed to blue light exhibit retrusion of the midface (arrowhead) and curved body not observed in the other groups. (F-Q) Alcian blue staining of cartilage and microdissection of the palate of 72 hpf embryos reveals a midface retrusion and cleft phenotype through the medial ethmoid plate (panel Q arrowhead) in the C120-*irf6*-ENR injected embryos grown under blue light (O-Q) which is not seen in control injected embryos (I-K) or injected embryos grown in the dark (L-N). Scale bars: 150um.

### Compound homozygote of *esrp1* and *esrp2* exhibits cleft lip and palate in zebrafish

To investigate the genetic requirement of *esrp1* and *esrp2* on zebrafish craniofacial development, CRISPR/Cas9 genome editing was utilized to generate *esrp1* and *esrp2* mutant alleles. Several alleles of *esrp1* and *esrp2* were were obtained, where alleles harboring −4bp and −14bp indels that lead to frameshift mutations and early protein truncation were selected for breeding, hereafter referred to as *esrp1*^−4bp/−4bp^ and *esrp2*^−14bp/−14bp^, respectively (Fig.S4). No phenotype was observed in the *esrp1*^−4bp/−4bp^ embryos, and *esrp2*^−14bp/−14bp^ fish developed normally except that females were infertile, as previously published in independently derived CRISPR alleles of *esrp1/2* (Burguera et al., 2017). However, compound homozygote *esrp1*^−4bp/−4bp^;*esrp2*^−14bp/−14bp^ zebrafish exhibit several phenotypes, consistent with previously published mutants (Burguera et al., 2017). The *espr1*^−4bp/−4bp^;*esrp2*^−14bp/−14bp^ embryos also failed to inflate the swim bladder, and the pectoral fins were formed but diminutive and the margins of the fin appeared dysplastic with irregular morphology. Further, Alcian blue staining of double knockouts revealed cleft in the anterior neurocranium, but the ventral cartilages, including the Meckel’s cartilage, were formed and appeared wild type (Fig. 5A).

**Figure 5.**
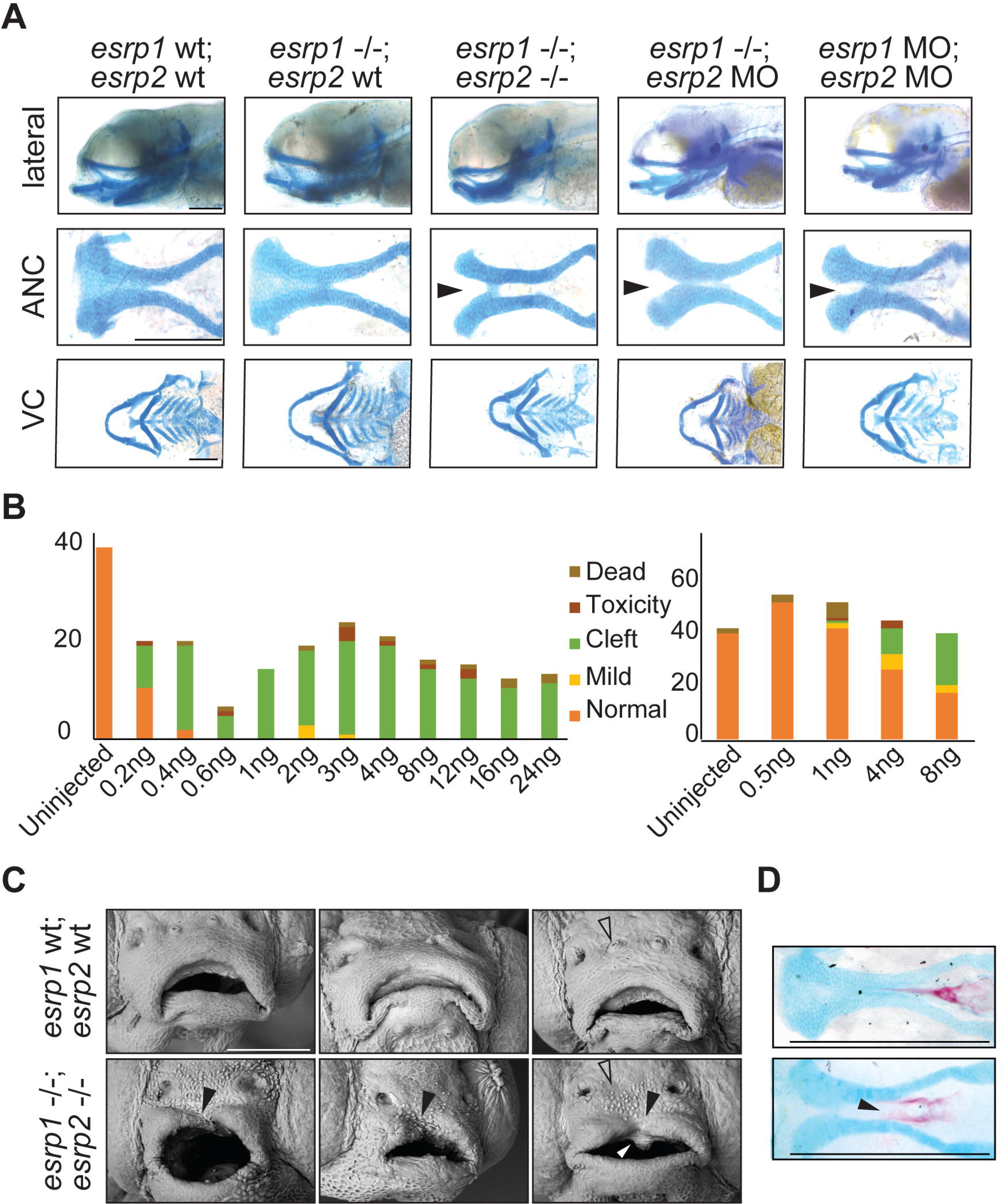
*esrp1/2* double mutants display a cleft lip and palate. (A) Alcian blue staining of 4 dpf zebrafish. Representative images of wild type, *esrp1* CRISPR mutant (*esrp1* −/−), and *esrp1/2* double CRISPR mutant (*esrp1* −/−; *esrp2* −/−), as well as *esrp1* CRISPR mutant treated with *esrp2* morpholino and wild type treated with *esrp1* and *esrp2* morpholino (*esrp1* MO, *esrp2* MO). Flat-mount images of the anterior neurocranium (ANC) show a cleft (arrow) between the median element and lateral element of the ANC when both *esrp1* and *esrp2* function were disrupted. Lateral images and flat-mount images of the ventral cartilage (VC) show only subtle changes in morphology between WT and *esrp1/2* −/− zebrafish. (B) Morphant phenotypes observed over a range of *esrp1* and *esrp2* morpholino doses. Single *esrp2* MO injections in the *esrp1*−/− background achieves nearly 100% phenotype penetrance, even at very low MO doses. (C) SEM of 5 dpf zebrafish showing discontinuous upper lip (closed arrow) in the *esrp1/2* double CRISPR mutant as well as absent preoptic cranial neuromasts (open arrow) and abnormal keratinocyte morphology. White arrow depicts aberrant cell mass (D) Representative images of alizarin red/Alcian blue staining of 9 dpf *esrp1/2* double CRISPR mutant zebrafish and wild type clutch-mate controls. *Esrp1/2* ablation causes abnormal morphology of the mineralizing parasphenoid bone, where the bone appears wider and with a cleft (arrow).

Inter-cross of *esrp1*^−4bp/−4bp^; *esrp2*^wt/−14bp^ produces predicted Mendelian ratio of 25% *esrp1*^−4bp/−4bp^; *esrp2*^−14bp/−14bp^ embryos for downstream phenotypic analysis, where 75% of the embryos appeared wild type. In order to increase the percentage of embryos that can be useful for analysis to 100%, we asked whether morpholino disruption of *esrp2* in the *esrp1*^−4bp/−4bp^ background would yield consistent a cleft ANC phenotype that phenocopied *esrp1*^−4bp/−4bp^; *esrp2*^−14bp/−14bp^ mutant. In fact, we were successful in phenocopying the cleft ANC phenotype by co-injecting *esrp1* and *esrp2* MOs into wild type embryos. However, the morpholino concentration needed were relatively high, requiring 2-8 ng of each MO to be injected for ~25-50% of embryos to develop a cleft (Fig. 5A,B). However, when *esrp1*^−4bp/−4bp^ embryos were injected with *esrp2* MO, the cleft ANC phenotype was consistent and observed in nearly 100% of the embryos, even when the MO concentration was reduced as low as 0.4 ng (Fig. 5A,B). One explanation for this phenomenon is the possibility that transcriptional compensation between *esrp1* and *esrp2* occurs when each gene is targeted, thereby requiring higher doses of each MO to ablate *esrp* activity sufficiently (Rossi, 2015 #1039). But when one of the *esrp* genes is already disrupted in the homozygous *esrp1*^−4bp/−4bp^ mutant, the threshold for full *esrp* loss of function is lower and requires\d a much smaller dose of MO to generate the cleft ANC phenotype.

Using scanning electron microscopy (SEM), one can observe that a cleft of the upper margin of the stomodeum invaginates and extends into the cleft ANC. Additionally the keratinocyte morphology of the surface epithelium appeared irregular and round with epithelial blebs in the *esrp1*^−4bp/−4bp^;*esrp2*^−14bp/−14bp^ embryo. In contrast, the wild type surface epithelium keratinocytes appeared octogonal or hexagonal without epithelial belbs (Fig. 5C). Alizarin red staining of the larvae at 9 dpf also revealed a lack of mineralization at the midline of the parasphenoid bone (Fig. 5D), consistent with a cleft ANC that remained clefted to ossification stage and subsequent larval fish development.

### Zebrafish ANC morphogenesis is dependent on epithelial interactions with infiltrating cranial neural crest cells

Formation of the zebrafish ANC involves migration of anteromost CNCCs to populate the median portion (frontonasal derived) while more posterior CNCCs migrate from each side (maxillary derived), these 3 discrete elements fuse to form the ANC. Concurrent with these cellular movements, the CNCCs undergo differentiation to chondrocytes (Dougherty et al., 2013; Wada et al., 2005). We found that the ablation of *irf6* (a key periderm/epithelial gene) and *esrp1/2* (epithelial-restricted genes) both resulted in a cleft in the ANC, where condrocytes are absent along the fusion plane between the frontonasal-derived median element and one side of the maxillary-derived lateral element (Figs. 4C and 5A).

To investigate the absence of these ANC chondrocytes, we performed lineage tracing of CNCCs in *esrp1/2* ablated embryos. Previously, we and others identified that the anteromost cranial neural crest populations at 20 somites that will migrates to and populates the median (frontonasal) element of the ANC (Dougherty et al., 2013; Wada et al., 2005). Accordingly, we labeled the CNCCs at 20-somite stage destined for the median element through photo-conversion of kaede under the lineage specificity of the *sox10* promoter. CNCCs of wild type or *esrp1/2* CRISPR mutants or *esrp1/2* morphants were photo-converted at 12-15 hpf (Fig. 6A,B). Embryos were imaged at 4 dpf to determine the population of the ANC contributed by photo-converted cells. We found that *esrp1/2* ablation did not affect the ability of CNCCs to migrate into the ANC, and reached posterior positions, without clustering anteriorly (Fig. 6A,B). Therefore, the cleft of the ANC in the *esrp1/2* mutants is not due to an absence of progenitor cells or defect in CNCC migration into the ANC. Nevertheless, Alcian blue staining confirmed that chondrocytes were absent from a cleft in the ANC in the *esrp1/2* mutants (Fig. 5A).

**Figure 6.**
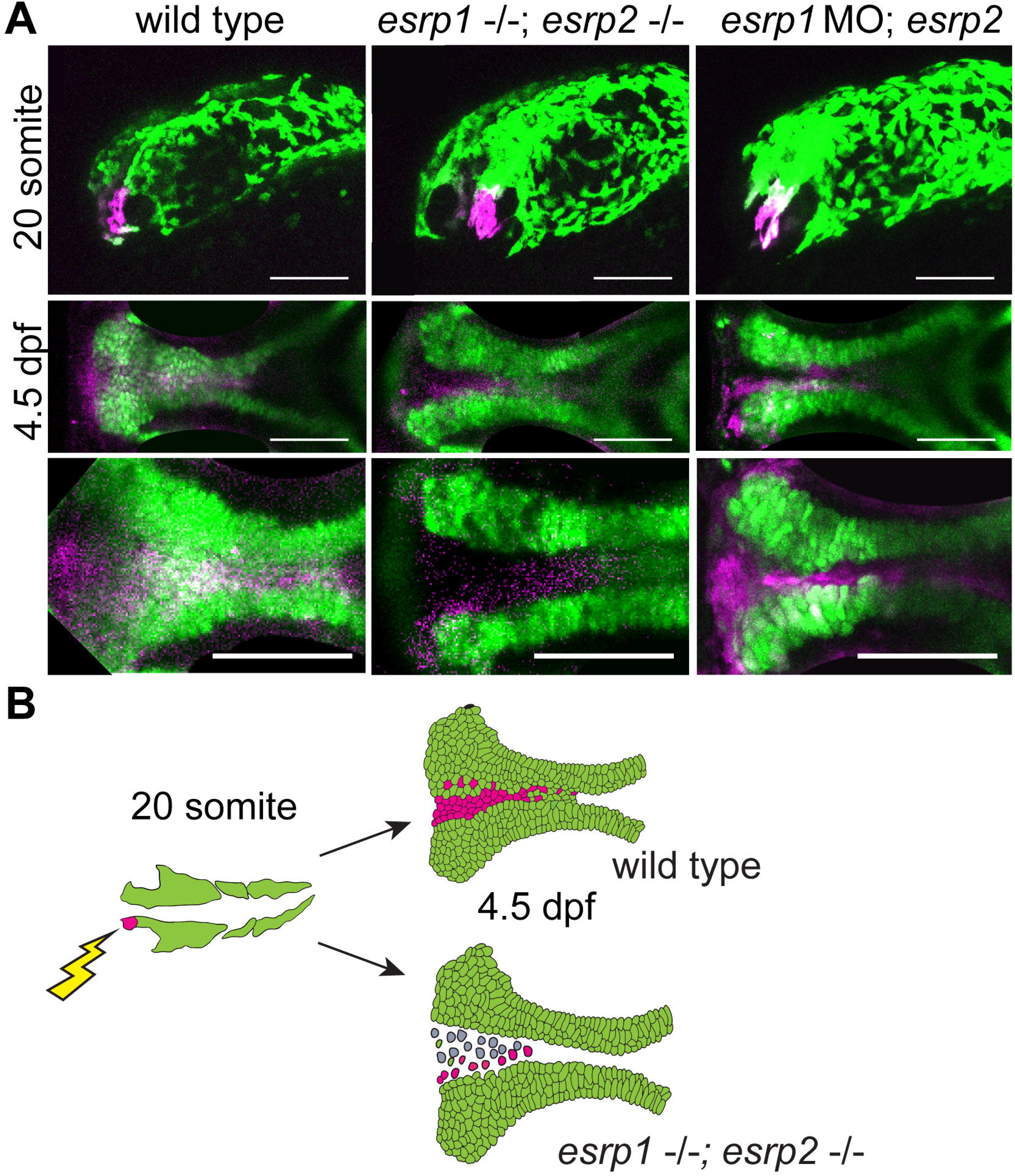
*esrp1/2*-null cranial neural crest cells migrate to the anterior neurocranium but do not differentiate to chondrocytes. (A) Lineage tracing of wild type or *esrp1/2* morphant zebrafish embryos using the Tg(sox10:kaede) line, native kaede fluorescence is shown in green, and photo-converted kaede is shown in magenta. Sagittal and horizontal views of zebrafish embryos at 19 hpf and 4.5 dpf, respectively. The anterior-most neural crest frontonasal prominence (FNP) progenitors were photoconverted at 19 hpf. At 4.5 dpf, the wild type signal tracks to the medial portion of the anterior neurocranium (ANC). Both the *esrp1*/esrp2 double CRISPR mutants and *esrp1/2* morphants exhibit a cleft in the ANC with absence of a portion of *sox10*-positive cells in the medial portion of the ANC, but the labeled CNCC representing FNP progenitors did reach and populate the entire length of the ANC. (B) Illustrative summary of lineage tracing results showing that photo-converted anterior most CNCCs contributing to FNP do migrate into the ANC in *esrp1/2* mutant embryos but a cleft forms at the juxtaposition of the FNP-derived median element and the maxillary-derived lateral element.

To investigate the cellular composition of the ANC cleft, we performed RNAscope ISH staining of coronal cryosections of wild type and *esrp1/ 2* mutants at 4 dpf. Sections through ANC clefts showed a dense population of cells in the location of the ANC cleft (Fig. 7A,B). In fact, this mass of cells can be localized in the SEM image of the *esrp1/2* mutant larvae (Fig. 5C). These cells are *col2a1* negative, consistent with absent Alcian blue staining. Instead, this abberant cell population in the position of the ANC cleft does express *irf6* and *krt4* (Fig. 7A,B). Confocal images of 5 dpf embryos confirms the expression of *irf6* and revealed *sox10* expression in these abberant cells (Fig. 8A). The expression of *sox10* suggests that at least a portion of these cells are CNC-derived whereas *krt4* expression indicates epithelial differentiation. The presence of *irf6* expression may be indicative of epithelial/periderm cells, or indicative of expression by CNCCs as has been previously reported (Dougherty et al., 2013; Kousa et al., 2019). Based on these results, we hypothesize that epithelial (or possibly periderm) cells that underly the frontonasal and maxillary prominence derivatives are defective in the *esrp1/esrp2* null mutants and prevented fusion of the median and lateral elements of the ANC, causing a cleft to form (Fig. 8B). In this way, this is the first direct evidence of cleft pathogenesis in the zebrafish due to epithelial defect, and forms a model to consider conservation of cleft pathogenesis involving the primary palate across vertebrates.

**Figure 7.**
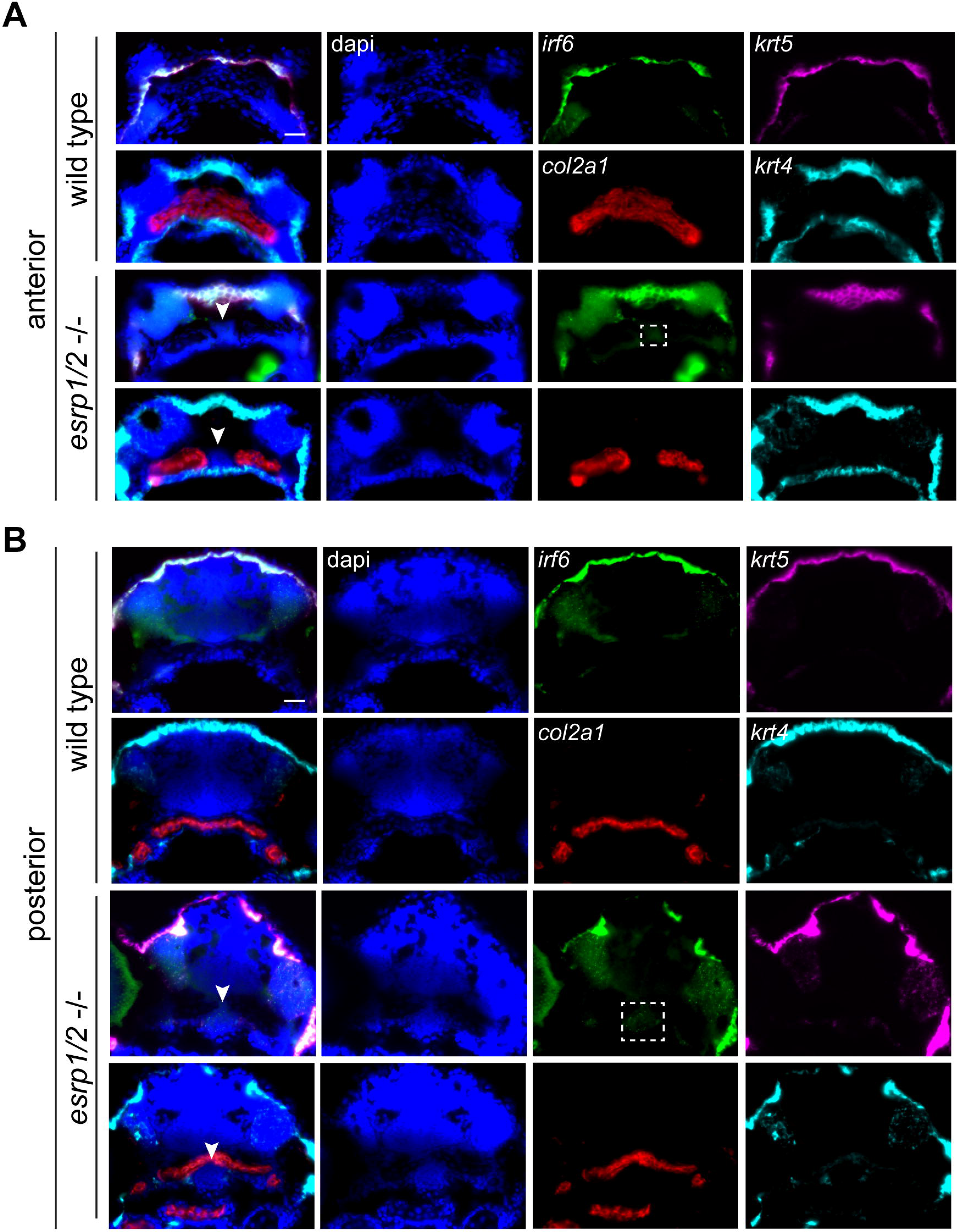
Anterior neurocranium of *esrp1/2* double mutants is populated by undifferentiated cells. Representative images of RNAscope ISH of coronal sections of *esrp1/2* double CRISPR mutants and wild type clutch-mate controls at 4 dpf. (A) Sections through ANC anterior to the eyes. *col2a1* (red) staining depicts normal morphology of the ANC cartilage elements in wild type while a cleft is apparent in the *esrp1/2*−/− zebrafish, with dapi (blue) stained cells between adjacent trabeculae (white arrow). These *col2a1* negative cells do not express epithelial markers *krt4* (cyan) or *krt5* (magenta), however there is low expression of *irf6* (green; white box). (B) Sections posterior to those in (A) show *col2a1* negative cells continuing inferior to the trabeculae in the *esrp1/2* mutant zebrafish and cells have low expression of *irf6* and *krt4* (white box). Scale bars: 20 μm.

**Figure 8.**
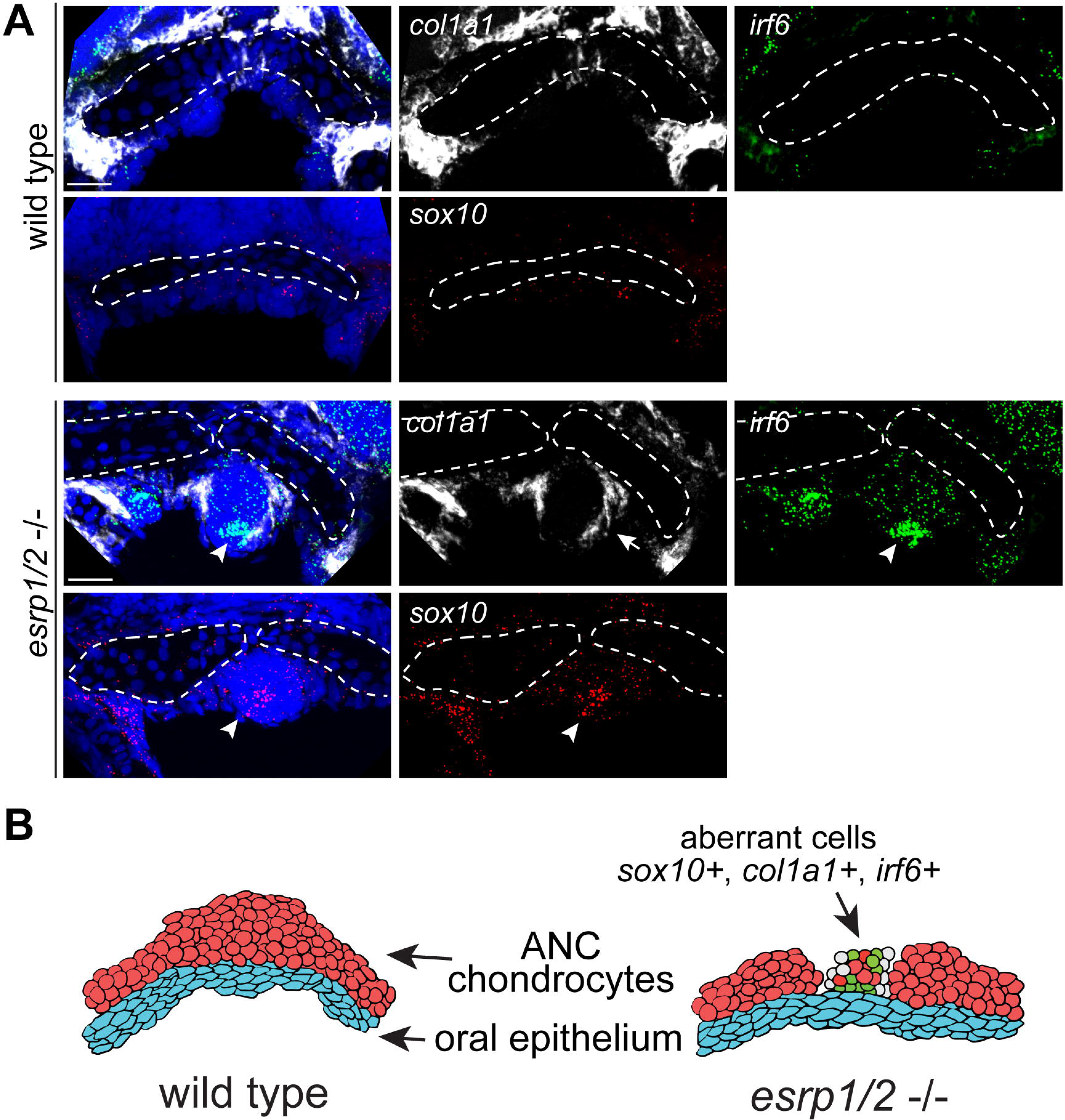
Aberrant anterior neurocranium cells of *esrp1/2* double mutants express CNCC and epithelial cell markers. Representative z-stacks of RNAscope ISH of coronal sections of *esrp1/2* double CRISPR mutants and wild type clutch-mate controls at 5 dpf. Sections through ANC anterior to the eyes. ANC cartilage elements are outlined in white. *col1a1* (white) staining depicts perichondrium surrounding the abberant mass of cells in the *esrp1/2* mutant zebrafish consistant with chondrogenic condensation (arrow). *Irf6* (green) and *sox10* (red) expression is apparent in these cells (arrow head). Dapi (blue). Scale bar: 20 μm. (B) Illustrative summary of RNAscope ISH results showing a heterogenous mix of cells populating the cleft between ANC cartilage elements.

### Histological analysis and comparison of *Irf6*^*R84C*^ and *Esrp1/2* knockout mice

*Irf6* and *Esrp1/2* are established epithelial-expressed genes, and their essential roles in the fusion of craniofacial processes have been reported (Bebee et al., 2015; Ingraham et al., 2006; Iwata et al., 2013). Interestingly, the phenotype of the *Irf6*−/− and *Irf6*^*R84C*^ mouse versus the *Esrp1/2*^−/−^ mouse differ significantly. Whereas *Esrp1/2*^*−/−*^ mice have a clear bilateral cleft lip and primary palate, *Irf6*^*−/−*^ mice do not exhibit a cleft in the lip. Previous analysis of the *Irf6*^−/−^ and *Irf6*^*R84C*^ mouse mutants reported a cleft of the secondary palate and a hypoplastic midface, which was attributed to impaired epithelial development (Ingraham et al., 2006; Richardson et al., 2006). However, as we examined the *Irf6*^R84C^ mutant embryos by histological staining of coronal sections, we found that the lip and premaxilla of the mutant embryo were hypoplastic beyond epithelial associated tissue, where the lip and facial muscles were significantly atrophic, the alveolar bone was deficient, and the incisor roots were displaced (Fig. 9). In contrast, histological analysis of the *Esrp1/2* double knockout shows clear development of epithelial and mesenchymal derived tissue with a bilateral lack of fusion between the medial and lateral prominences of the midface (Fig. 9). Therefore, while the *Irf6*^*R84C*^ mutant did not exhibit a cleft of the lip or the primary palate, the midfacial developmental phenotype is more severe.

**Figure 9.**
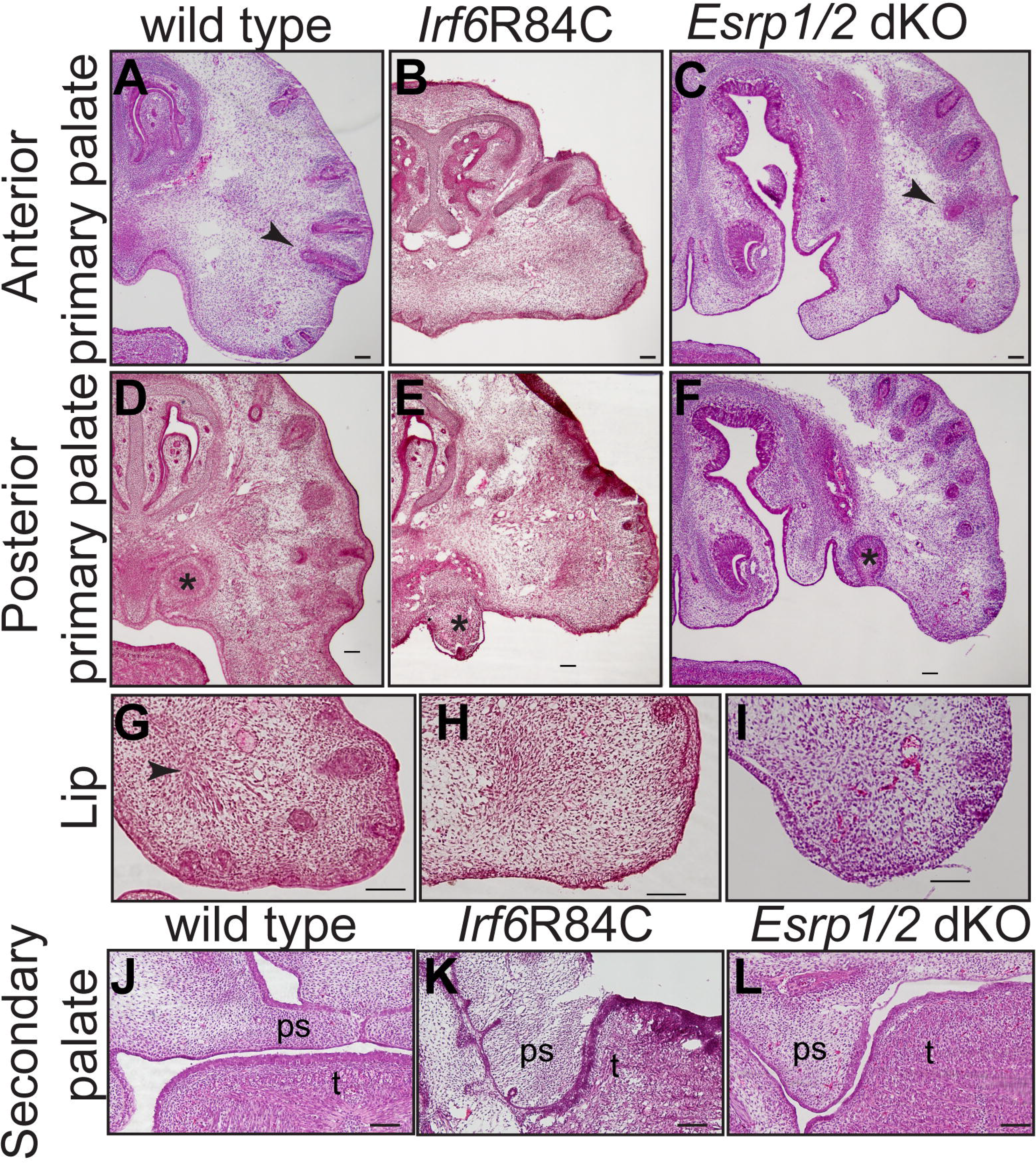
*Irf6*^*R84C/R84C*^ and *Esrp1/2* dKO mice display different cleft characteristics. Hematoxylin and eosin staining of coronal cryosections of wild type, *Irf6*^*R84C/R84C*^ and *Esrp1/2* dKO. (A-F) E13.5 asterisk: tooth primordium, closed arrowhead: muscle, open arrowhead: vibrissae. (G-I) Magnified images of E13.5 lip tissue showing differences in tissue composition. (J-L) E15.5 sections through secondary palate shelves. Both *Irf6*^*R84C/R84C*^ and *Esrp1/2* dKO mice exhibit a secondary cleft with impaired elevation of the palatal shelves. However, whereas failed elevation in *Irf6*^*R84C/R84C*^ mice has been attributed to eplithelial adhesions, *Esrp1/2* dKO mice do not exhibit adhesion. Scale bars: 100 μm.

### Genetic interaction of *Irf6R84C* with *Esrp1* and *Esrp2*

*Irf6*^*R84C*^ heterozygous mice have some epithelial adhesions; however, these adhesions are reported to resolve and heterozygous mice develop normally (Ingraham et al., 2006; Richardson et al., 2006). *Esrp1* heterozygotes also develop normally, and *Esrp2* hetero and homozygous mice have no phenotype. However, the compound ablation of *Esrp1* with *Esrp2* worsens the cleft lip phenotype observed is the *Esrp1* mutant (Bebee et al., 2015).

To test the hypothesis that *Irf6* and *Esrp1/2* function in the same developmental pathway, we carried out genetic epistasis analysis and generated *Irf6*;*Esrp1*;*Esrp2* compound mutants. We hypothesized that if *Irf6* and *Esrp1/2* genetically interacted, that *Irf6* and *Esrp1* heterozygosity on an *Esrp2* null background may result in a cleft phenotype, when *Irf6* and *Esrp1* heterozygotes do not normally form a cleft. As expected, we observed that *Irf6*^*R84C/+*^;*Esrp1*^+/−^;*Esrp2*^+/−^ mice developed and reproduced normally. To generate *Irf6*^*R84C/+*^;*Esrp1*^+/−^;*Esrp2*^−/−^ embryos, we intercrossed the triple heterozygous mice. We collected a total of 49 embryos from 7 litters from E12.5-E18.5 and tabulated the resulting genotypes (Table 1; Table S1). Based on Mendelian genetics, we expected approximately 3 *Irf6*^*R84C/+*^;*Esrp1*^+/−^;*Esrp2*^−/−^ embryos; however, no triple homozygous mutant embryos were collected (Table 1) suggesting that the genotype is prenatally lethal. Additionally, we did not collect any *Irf6*^*R84C/R84C*^;*Esrp1*^−/−^ embryos, which indicates this genotype is also lethal (Table 1). These findings suggest cummulative lethality between *Irf6* and *Esrp1/2,* where non-lethal genotypes produce a lethal phenotype when combined in the compound mutant.

**Table 1.** Irf6, Esrp1 and Esrp2 genotypes interact to produce non-Mendelian embryo ratios.

| *Irf6* | *Esrp1* | *Esrp2* | probability | Expected | Observed |
| --- | --- | --- | --- | --- | --- |
| Het | WT | WT | 1/32 | 1.5 | 3 |
| Het | WT | Het | 1/16 | 3.1 | 4 |
| Het | WT | KO | 1/32 | 1.5 | 1 |
| Het | Het | WT | 1/16 | 3.1 | 7 |
| Het | Het | Het | 1/8 | 6.1 | 3 |
| **Het** | **Het** | **KO** | **1/16** | **3.1** | **0** |
| Het | KO | WT | 1/32 | 1.5 | 1 |
| Het | KO | Het | 1/16 | 3.1 | 5 |
| Het | KO | KO | 1/32 | 1.5 | 0 |
| R84C | WT | WT | 1/64 | 0.8 | 0 |
| R84C | WT | Het | 1/32 | 1.5 | 2 |
| R84C | WT | KO | 1/64 | 0.8 | 0 |
| R84C | Het | WT | 1/32 | 1.5 | 3 |
| R84C | Het | Het | 1/16 | 3.1 | 3 |
| R84C | Het | KO | 1/32 | 1.5 | 3 |
| R84C | KO | WT | 1/64 | 0.8 | 0 |
| R84C | KO | Het | 1/32 | 1.5 | 0 |
| R84C | KO | KO | 1/64 | 0.8 | 0 |

*Irf6*<sup>R84C/WT</sup>; *Esrp1*<sup>+/-</sup>; *Esrp2*<sup>+/-</sup> triple heterozygous mice were in-crossed and embryos were collected between E12.5 and E21. Table 1 is a subset of expected number of embryos based on Mendelian genetics versus the observed number of viable embryos. The *Irf6*<sup>R84C/WT</sup>; *Esrp1*<sup>+/-</sup>; *Esrp2*<sup>-/-</sup> genotype appears to be lethal prior to E12.5 as approximately 3 embryos were expected but zero embryos were observed. A total of 49 embryos were collected from 7 different litters.

## DISCUSSION

Orofacial clefts are a common birth defect, and GWASs have identified some critical genes associated with syndromic and non-syndromic cleft. Here we described mouse and zebrafish models using genes with known genetic variants in human cleft patients, *IRF6* and *ESRP1/2*. We present evidence to support that *Irf6* and *Esrp* function in the same regulatory pathway. We observed that mz-*irf6*^−8bp/−8bp^ zebrafish embryos have significantly decreased expression of *esrp1* and this is rescued upon introduction of *irf6* mRNA. This finding is consistant with *esrp1* being a transcriptional target of *irf6* and putative *irf6* response elements (Khan et al., 2018) can be found surrounding the *esrp1* transcriptional start site. Additionally, a RNA-seq experiment identified known *irf6* targets including *grhl3* and *tfap2a*. Direct molecular experiments are needed however to test transcriptional regulation of *esrp1* by *irf6*. We found that *Irf6* and *Esrp1/2* are consistently co-expressed in the embryonic frontonasal ectoderm and oral epithelium associated with the palate, and epithelium of the mouth opening, in both mouse and zebrafish.

In zebrafish, *irf6* null embryos ruptured during gastrulation whereas *esrp1/2* null embryos survive to larval stage. However, post-gastrulation ablation of *irf6* resulted in a similar cleft morphology of the ANC as the *esrp1/2* null. Further analysis of the *esrp1/2* null showed that the cleft of the ANC correlated with a cleft in the upper margin of the mouth opening, reminiscent of a human cleft lip. Further, using an NCC-specific photo-convertible reporter line, we were able to show that migration of CNCC to the developing ANC occured but chondrogenesis was impaired.

This work highlights the utility of complementary studies of palate morphogenesis in zebrafish and mouse models. The zebrafish model affords the transgenic tractability and visualization of CNCC migration, enabling us to determine the cellular mechanism responsible for the cleft ANC. The mouse mutants provide the mammalian anatomic contexts to examine cleft malformation. *Irf6*^*R84C*^ mice have significant midfacial hypoplasia and oral occlusion when compared to *Esrp1/2* null embryos that exhibit bilateral cleft lip and palate. Histological examination of *Irf6*^*R84C*^ mice revealed loss of portions of normal mesenchymal-derived tissue (muscle, nerve, bone, and vascularity). In contrast, the mesenchymal derived tissues are fully formed but displaced in the *Esrp1/2* null mice.

The early lethality of *irf6* null zebrafish at the onset of gastrulation initially precluded analysis of the *irf6* zygotic requirement in craniofacial development. Here, we utilized an optogenetics strategy to disrupt *irf6* function after gastrulation when the embryonic body axis had formed, thereby revealing the zygotic requirement for *irf6*. Future studies will use the *irf6* optogenetic model to study the roles of *irf6* during ANC and lip morphogenesis. Interestingly, periderm markers identified in the mouse lambdoidal junction were found to be dysregulated in the *irf6* mutant zebrafish model, specifically *grhl3*, *tfap2a* and *perp*. Additionally *gata3,* which was identified as a mesenchymal marker at the fusion zone of mice (Li et al., 2019), is dysregulated in the irf6 null zebrafish.

While the *esrp1*^*−4bp/−4bp*^*;esrp2*^*−14bp/−14bp*^ zebrafish exhibited a consistent cleft lip and palate phenotype, the infertility of the *esrp2*^*−14bp/−14bp*^ fish preclude large-scale experiments to analyze downstream mechanisms of the development of cleft palate. We generated a robust *esrp1*^*−4bp/−4bp*^; *esrp2* morphant assay that can be applied to toward chemical screening experiments and functional testing of human *ESRP1/2* gene variants.

In humans, CPO is less common than CL/P (Bush and Jiang, 2012; Gritli-Linde, 2008; Van Otterloo et al., 2016). Although humans and mice share approximately 99% of their genes and the early craniofacial development of the mouse embryo closely mirrors the human, including gene expression (Swartz et al., 2011), there is a striking difference in the manifestation of orofacial cleft defects (Gritli-Linde, 2008). Most often when a human CL/P-associated gene has been disrupted in mice, a cleft of the palate forms but the lip appears normal. Our current understanding in humans is that CL/P and CPO are different genetic disorders (Dixon et al., 2011; Gritli-Linde, 2008; Juriloff and Harris, 2008). These discrepencies between humans and mouse models hampens understanding of the etiopathogenesis of human CL/P. Here we characterize the *Esrp*1/2 null mouse, exhibiting bilateral CL/P, as an important model for studying OFC etiopathogenesis. Additionally, as we place *ESRP1* in the *IRF6* gene regulatory pathway, we hope to better understand how alternative isoforms regulated by *ESRP1* may in turn be important for palate development.

Whereas zebrafish have historically been an excellent model organism for forward genetic screens, CRISPR/Cas9 gene editing technology has permitted relatively efficient reverse genetic engineering of zebrafish (Liu et al., 2019). This utility of the zebrafish embryo for studying developmental processes and modeling human cleft-associated genes necessitates further study into their craniofacial morphogenesis. Transplant and lineage tracing experiments have illuminated the neural crest origin of the zebrafish ANC, and how the frontonasal and paired maxillary cartilage elements converge into a continuous cartilage structure (Dougherty et al., 2013; Dougherty et al., 2012; Wada et al., 2005). We show that *IRF6* and *ESRP1* are conserved in their requirement for ANC morphogenesis, where disruption result in orofacial cleft in human, mouse and zebrafish. These findings provide evidence of conserved molecular and morphological processes occurring in the merging and fusion of the mouse and zebrafish midface.

We suspect that non-epithelial expression of *Irf6* contributes to normal craniofacial morphogenesis and may explain some differences in the *Irf6* and *Esrp* mutant phenotypes. Future research utilizing tissue-specific knockout of *Irf6* will address this hypothesis. We also suspect that the *Irf6* phenotype is more severe because *Irf6* acts upstream of *Esrp1*, along with additional targets, and ongoing experiments on the transcriptional activity of *Irf6* will be important. Continued breeding of the triple heterozygous *Irf6*^*R884C/+*^, *Esrp1*^+/−^; *Esrp2*^+/−^ mice will also test a genetic epistasis interaction between them. Recently an in-depth analysis of a lineage-specific *Esrp1* knockout mouse was completed and found that *Esrp1* regulates proliferation of the mesenchyme of the lateral nasal prominences, along with being required for fusion of the medial and lateral nasal prominences (Lee et al., 2020). The *Esrp1/2* knockout mouse is an exciting model for studying craniofacial development and the etiology of CL/P, as relatively few mouse models exhibit cleft lip phenotype. Ongoing work to identify *Esrp1/2* molecular targets and mechanistic studies of these targets will provide new insight into palate morphogenesis.

## METHODS

### Animal breeding and gene editing

All animal experiments were performed in accordance with protocols approved by Massachusetts General Hospital Animal Care and Usage Committee. C57Bl/6J (wild type) animals were obtained from the Jackson Laboratory. *Irf6*^*R84C/+*^ mice were received as a gift from Dr. Yang Chai. *Esrp1*^*+/−*^*;Esrp2*^*−/−*^ mice were received from Dr. Russ Carstens. Embryonic day 0.5 was considered to be noon on the day of the copulatory plug.

Zebrafish (*Danio rerio*) adults and embryos were maintained in accordance with approved institutional protocols at Massachusetts General Hospital. Embryos were raised at 28.5°C in E3 medium (5.0 mM NaCl, 0.17 mM KCl, 0.33 mM CaCl_2_, 0.33 mM MgSO_4_) with 0.0001% methylene blue. Embryos were staged according to standardized developmental timepoints by hours or days post fertilization (hpf or dpf, respectively).(Kimmel et al., 1995) All zebrafish lines used for experimentation were generated from the Tübingen strain.

CRISPR sgRNA target sites were identified by a variety of online CRISPR computational programs such as zifit.partners.org/ZiFiT(Sander et al., 2007), crispr.mit.edu (Ran et al., 2013), and chopchop.rc.fas.harvard.edu.(Montague et al., 2014) sgRNAs were designed with the traditional sequence constraint of a 3’ PAM sequence containing NGG and an additional sequence constraint of a 5’ NG for *in vitro* RNA synthesis.

The *esrp1, esrp2* and *irf6* CRISPR sgRNAs were generated by *in vitro* transcription from a SP6 promoter as described.(Gagnon et al., 2014) Lyophilized Cas9 protein (PNA Bio) was resuspended in ddH_2_O to a stock concentration of 1 μg/μl and stored in single-use aliquots in −80°C to avoid freeze-thaw inactivation and kept for 6 months. One-cell staged zebrafish embryos were microinjected directly in the cytoplasm with 2 nl of a solution containing 15 ng/μl of sgRNA and 100 ng/μl of Cas9 protein pre-complexed for 5-10 minutes at room temperature. A subset of embryos injected with the sgRNA and Cas9 protein mixture was harvested for genomic DNA to confirm the presence of indels, and the rest were grown into adulthood as F0 mosaic fish. F0 adult fish were subsequently outcrossed with wild-type fish to generate F1 founders with germline transmission of indel alleles. F1 founders were further outcrossed with wild-type fish to yield a large number of heterozygote fish, and minimize the presence of off-target edits. Lastly, F2 heterozygotes were incrossed to generate homozygote embryos for phenotypic analysis.

Genomic DNA for genotyping was isolated from either whole 24 hpf embryos or tail fin clips using the HotSHOT method as described.(Meeker et al., 2007) Genotyping primers flanking the CRISPR sgRNA site were designed using a combination of ChopChop (chopchop.rc.fas.harvard.edu) and NCBI primer BLAST (ncbi.nlm.nih.gov/tools/primer-blast/). Forward primers were synthesized by Invitrogen with 5’-FAM modifications. Microsatellite sequencing analyses were used to determine indel mutation sizes and frequencies (MGH DNA Core), and Sanger sequencing was performed on PCR amplicons of CRISPR sgRNA to confirm the exact sequence changes resulting from CRISPR mutagenesis.

### mRNA sequencing and qPCR

Total RNA was isolated from 4 hpf wild type and maternal-null *Irf6*^*−/−*^ embryos by TRIzol and phenol-chloroform ethanol precipitation, as previously described. Total RNA was quantified with the Nanodrop 2500 and assessed for quality with Bioanalyzer 2100 RNA chips (Agilent). Samples with RNA integrity numbers (RIN) over 9 were selected to proceed with sequencing library preparation. mRNA-seq libraries were prepared with the NEBNext Ultra RNA library preparation kit with poly(A) mRNA magnetic isolation module (NEB) essentially according to manufacturer protocols. Resulting cDNA libraries were quantified by a Qubit fluorometer and assessed for quality with a Bioanalyzer. The sequencing-ready cDNA libraries were quantified with the NEBNext library quantification kit for Illumina (NEB). mRNA-seq libraries were sequenced with single-end 50 at ≈20 million reads per sample with biological triplicates. For qPCR, ~ 30 embryos per sample were collected in RLT buffer (Qiagen) and flash-frozen in liquid nitrogen. Samples were homogenized using a rotor-stator homogenizer, and RNA was isolated using an RNeasy Mini Kit (Qiagen). Total mRNA was quantified using a Nanodrop spectrophotometer and used for cDNA synthesis (Thermo). qPCR was performed with Taqman probes and reagents (Thermo), and expression was normalized to 18s rRNA expression.

### Zebrafish embryo microinjection of mRNA and morpholinos

Microinjection of mRNA was performed by injecting 2 nl of mRNA solution with 0.05% phenol red directly into the cytoplasm of one-cell staged embryos. Lyophilized morpholinos were resuspended with ddH2O to a stock concentration of 20 ng/μl and stored at RT in aliquots. Individual aliquots were heated to 70°C and briefly vortexed before preparation of the injection mix to ensure full dissolution. Mismatch control morpholinos were injected under identical conditions to control for potential toxicities. Embryos from all methods of microinjection were examined at 3 hpf to remove unfertilized embryos, which were quantified against the total number of microinjected embryos to ensure no fertilization defects were observed.

### Whole-mount *in situ* hybridization

Embryos were isolated at various time points and fixed in 4% formaldehyde at 4°C for 12-16 hrs. Subsequently, embryos were washed and stored in methanol. WISH and DIG-labeled riboprobes were synthesized as described.(Thisse and Thisse, 2008) Briefly, for riboprobe synthesis, PCR was performed using embryonic cDNA as templates and T7 promoter sequence-linked reverse primers to generate cDNA templates for *in vitro* transcription. PCR reactions were purified using the NucleoSpin gel and PCR clean-up kit (Machery-Nagel). *In vitro* transcription was performed using a T7 polymerase (Roche) and DIG labeling mix (Roche). DIG-labeled riboprobes were isolated with ethanol-NaOAc precipitation, resuspended in DEPC-treated ddH_2_O, and stored at −20°C. All PCR products were TOPO cloned into pGEM-T Easy vectors (Promega) and sequence verified by Sanger sequencing. WISH colorimetric signal detection was performed using an alkaline phosphatase-conjugated anti-DIG antibody (Roche) and BM Purple AP substrate (Roche).

### RNAscope *in situ* hybridization

Zebrafish and mouse embryos were fixed in 4% formaldehyde, taken through a sucrose gradient and cryo-embedded and sectioned. Probes were designed and purchased from ACD Bio, and hybridization and staining was performed according to the manufacturer’s protocol. Stained sections were imaged using either a confocal microscope, where a z-stack was obtained and analyzed on ImageJ for z-stack maximum intensity projections or a standard fluorescent microscope.

### Skeletal staining and brightfield imaging

Zebrafish embryos were fixed at 96 hpf or 120 hpf in 4% formaldehyde and stored at 4°C overnight, washed with PBS, dehydrated in 50% ethanol, and stained with acid-free Alcian blue overnight on a rotating platform at RT as described.(Thisse and Thisse, 2008) Stained embryos were washed with ddH2O and subsequently bleached (0.8% W/V KOH, 0.1% Tween 20, 0.9% H2O2) until cell pigmentation was no longer present. For double-stained embryos with Alcian blue and alizarin red, embryos were stained with a 0.05% alizarin red solution in ddH2O for 30 minutes on a rotating platform at RT following bleaching with KOH and H2O2. Afterward, double-stained embryos were placed in three changes of a tissue-clearing solution consisting of 25% glycerol and 0.1% KOH, each for 25 min. Whole and dissected stained embryos were mounted in 3% methylcellulose on a depression slide and imaged using a Nikon Eclipse 80i compound microscope with a Nikon DS Ri1 camera. Z-stacked images were taken to increase the depth-of-field with the NIS Element BR 3.2 software. Stacked images were processed by ImageJ to generate maximum intensity projection images.

For scanning electron microscopy (SEM), 4 dpf embryos were fixed in half-strength Karnovsky fixative. Samples were processed, and images were obtained by CBSET, Inc. Lexington, MA.

### Optogenetic expression of *irf6* in zebrafish

Genes *irf6*, *irf6*-ENR, i*rf6*^*R84C*^, and mCherry were isolated by PCR from various templates and inserted into the pGL4.23-(C120×5)-TATA vector with In-Fusion cloning (Clontech) according to manufacturer instructions using a 1:2 vector-to-insert ratio to generate optogenetic response plasmids. The constructs were transformed in Stellar chemically competent cells (Clontech), and colonies were screened by PCR, restriction digests, Sanger sequencing, and whole plasmid sequencing to verify the sequence identities and accuracy of the constructs. Light-sensitive response protein VP16-EL222 was subcloned into pCS2+8 and in vitro transcribed from the SP6 promoter as described above to generate capped mRNA for embryo microinjections. The optogenetics injection mix was comprised of 25 ng/μl EL222 and 10 ng/μl pGL4.23 response plasmid with 0.05% phenol red. Each embryo was microinjected with 2 nl of the optogenetics injection mix directly in the cytoplasm at the one-cell stage, immediately wrapped in aluminum foil, and placed into a dark incubator. Unfertilized and abnormal embryos were removed at 3 hpf in the dark room with limited exposure to ambient light. Injected embryos were divided into two groups (dark and light) at the desired developmental stage in E3 media without methylene blue and placed under 465nm blue light (LED panel, HQRP) at 0.3 mW/cm2 (measured by a PM100D digital power meter with an SV120VC photodiode power sensor, ThorLabs) with constant illumination. Control embryos containers were wrapped in aluminum foil.

### Lineage Tracing

Embryos originating from an *espr1*^−4bp/−4bp^;*esrp2*^+/−14bp^ incross, wild type embryos injected with 8ng esrp1 MO and 4ng esrp2 MO at the one-cell stage, or uninjected wild-type embryos, all in a Tg(*sox10:kaede*) background were grown until 20 somites, oriented for imaging in the sagittal position, and encased in 1% low-melt agarose. Using the 405nm UV laser and ROI setting in a Leica SP8 confocal microscope, the anterior-most portion of NCCs that contribute to the FNP were unilaterally photoconverted, keeping the alternate side as an internal control, as previously described.(Dougherty et al., 2012) Photoconverted embryos were carefully microdissected out of the agar, and grown in E3 at 28.5°C until 4dpf and imaged again to track the photoconverted cells. Maximum-projections of the photoconverted half of the embryo, or the planes consisting of the palate, in 14hpf or 4dpf embryos, respectively, were generated using Fiji/ImageJ.

## Acknowledgement

We ae grateful to Jessica Bethoney for excellent management of our aquatics facility. We thank Irimia Manuel for willingness to share an *esrp1/2* zebrafish mutant alleles (did not survive shipment, not used in this study) and Yang Chai for sharing the *Irf6*^*R84C*^ mouse mutant allele. The Esrp1 and Esrp2 mouse alleles were generated by R.C. with funding from the National Institutes of Health (R01DE024749). This work was supported by grants to E.C.L. from the National Institutes of Health (R01DE027983), the Shriners Hospital for Children, and the Laurie and Mason Tenaglia Massachusetts General Research Scholar Award.

