## Supplemental Figures for "Requirement of *Irf6* and *Esrp1/2* in frontonasal and palatal epithelium to regulate craniofacial and palate morphogenesis in mouse and zebrafish"

**Figure S1. Gene ontology (GO) gene-concept network analysis of RNA-seq data.** (A) *irf6*-/- embryos have perturbations in biological processes such as cell adhesion, gastrulation, mesoderm, ectoderm and endoderm development, and response to xenobiotic stimulus. (B) *irf6*-/- embryos also have perturbations of various cellular compartments including cell junctions, cytoskeletal elements and extracellular space. Grey nodes show GO terms, colored nodes show individual genes from the RNA-seq dataset, and edges connect genes to one or more associated GO terms. Colored nodes show relative enrichment (measured by fold-change) of genes in wild type samples relative to *irf6-/-* embryos. Maps were generated using the enrichplot package in R.

**Figure S2. GO Enrichment analysis of the wt vs. mz-*irf6*-/- RNA-seq dataset.** Nodes show GO terms associated with enriched genes in the dataset. Edges and arrows show previously annotated relationships between GO terms. Developmental processes such as ectodermal development and otic placode formation are enriched in the analysis, which involve the recruitment of neural crest cell progenitors also involved in palate development. Color represents q-value, with red having a lower q-value and blue having a higher q-value. Q-values were computed using FDR with the Benjamini-Hochberg correction. Data were visualized using the enrichGO and goplot packages in R.

**Figure S3. KEGG Graph of Cell Adhesion Molecules.** KEGG pathway analysis that maps enriched genes in our dataset to a database containing cellular pathway information. Colored nodes show increased expression of genes in red and decreased expression in green for wild type fish compared to control. Many cell-adhesion molecular interactions are disrupted in the *irf6*-/- embryos. Data were generated using the enrichKEGG and pathview packages in R.

**Figure S4. Generation of *esrp1* and *esrp2* CRISPR/Cas9 gene disruption in zebrafish.** (A) Schematic representations of *esrp1* and *esrp2* exons, positions of target site (arrow), and sequences of mutations. (B) Micro-satellite results of genotyping PCR showing a size shift consistent with the observed mutation. (C) Expression of the CRISPR/Cas9 targeted gene is decreased relative to wild type, as measured by qPCR. n=3,4. $p<0.01$

**Table S1. *Irf6*, *Esrp1* and *Esrp2* genotypes interact to produce non-Mendelian embryo ratios.** Expected number of embryos based on Mendelian genetics of *Irf6*$^{R84C/WT}$; *Esrp1*$^{+/-}$; *Esrp2*$^{+/-}$ triple heterozygous in-cross versus the observed number of viable embryos. *Irf6*$^{R84C/WT}$; *Esrp1*$^{+/-}$; *Esrp2*$^{+/-}$ triple heterozygous mice were in-crossed and embryos were collected between E12.5 and E21. A total of 49 embryos were collected from 7 different litters.

### Figure S1

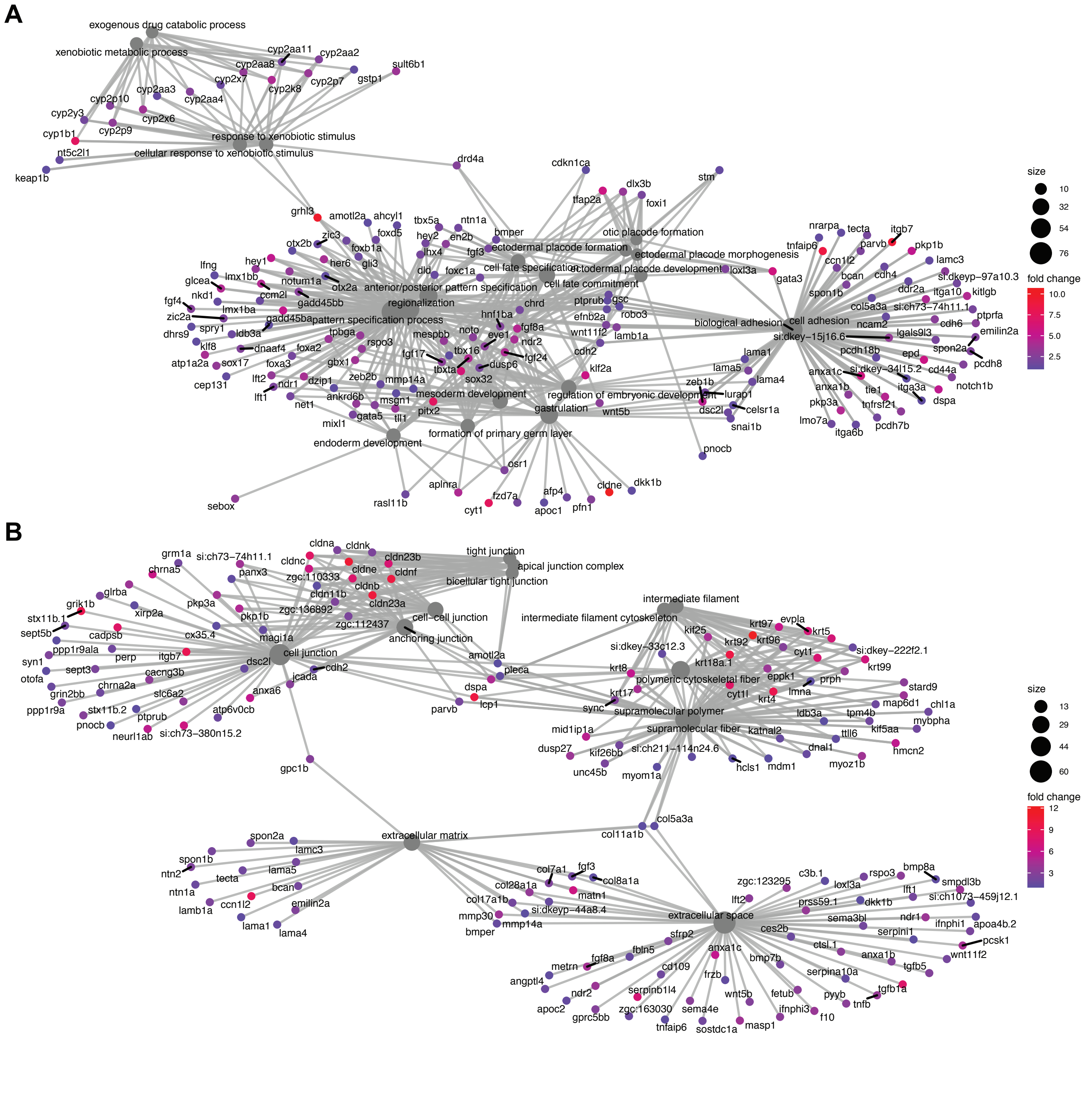
A
exogenous drug catabolic process
xenobiotic metabolic process
cyp2aa11
cyp2aa2
cyp2aa8
sult6b1
cyp2x7
cyp2p7
gstp1
cyp2k8
cyp2aa3
cyp2p10
cyp2aa4
cyp2y3
cyp2x6
cyp2p9
cyp1b1
response to xenobiotic stimulus
nt5c2l1
cellular response to xenobiotic stimulus
keap1b
drd4a
cdkn1ca
stm
dlx3b
tfap2a
foxi1
grhl3
amotl2a
ahcyl1
tbx5a
ntn1a
foxd5
otic placode formation
zic3
hey2
en2b
bmper
otx2b
foxb1a
lhx4
ectodermal placode formation
ectodermal placode morphogenesis
hey1
her6
gli3
fgf3
dld
foxc1a
cell fate specification
ectodermal placode development
loxl3a
gata3
lfng
lmx1bb
notum1a
otx2a
anterior/posterior pattern specification
cell fate commitment
glcea
ccm2l
nkd1
gadd45bb
regionalization
chrd
ptprub
gsc
fgf4
lmx1ba
robo3
zic2a
gadd45ba
pattern specification process
efnb2a
spry1
hnf1ba
fgf8a
biological adhesion
dhrs9
ldb3a
tpbga
noto
eve1
wnt11f2
lamb1a
klf8
dnaaf4
rspo3
mespbb
ndr2
atp1a2a
foxa2
sox17
gbx1
fgf17
tbx16
fgf24
foxa3
tbxta
dusp6
cdh2
lft2
zeb2b
sox32
klf2a
cep131
ndr1
dzip1
mmp14a
lft1
ankrd6b
mesoderm development
regulation of embryonic development
net1
msgn1
pitx2
gastrulation
wnt5b
mixl1
gata5
tll1
formation of primary germ layer
endoderm development
osr1
sebox
rasl11b
aplnra
fzd7a
cldne
dkk1b
afp4
cyt1
apoc1
pfn1
pnocb
nrarpa
tecta
itgb7
tnfaip6
parvb
pkp1b
ccn1l2
lamc3
bcan
cdh4
si:dkeyp−97a10.3
spon1b
ddr2a
itga10
kitlgb
col5a3a
si:ch73−74h11.1
ptprfa
ncam2
cdh6
cell adhesion
emilin2a
si:dkey−15j16.6
lgals9l3
pcdh18b
spon2a
lama1
epd
pcdh8
lama5
cd44a
anxa1c
si:dkey−34l15.2
notch1b
zeb1b
lama4
anxa1b
tie1
itga3a
lurap1
pkp3a
dspa
tnfrsf21
dsc2l
celsr1a
lmo7a
pcdh7b
itga6b
snai1b
size
10
32
54
76
fold change
10.0
7.5
5.0
2.5
B
grm1a
si:ch73−74h11.1
cldna
cldnk
tight junction
chrna5
cldnc
cldn23b
apical junction complex
panx3
cldne
cldnf
zgc:110333
bicellular tight junction
cldnb
glrba
cldn11b
pkp3a
cldn23a
grik1b
xirp2a
zgc:136892
pkp1b
cell−cell junction
stx11b.1
zgc:112437
cx35.4
sept5b
cadpsb
magi1a
anchoring junction
ppp1r9ala
perp
itgb7
cell junction
amotl2a
syn1
dsc2l
cdh2
sept3
cacng3b
pleca
otofa
jcada
chrna2a
dspa
grin2bb
slc6a2
anxa6
lcp1
ppp1r9a
stx11b.2
parvb
ptprub
atp6v0cb
pnocb
neurl1ab
si:ch73−380n15.2
gpc1b
intermediate filament
intermediate filament cytoskeleton
krt97
evpla
kif25
krt5
krt92
krt96
si:dkey−33c12.3
cyt1
si:dkey−222f2.1
krt18a.1
krt99
krt8
eppk1
polymeric cytoskeletal fiber
prph
stard9
krt17
cyt1l
lmna
krt4
map6d1
chl1a
sync
supramolecular polymer
ldb3a
tpm4b
mybpha
mid1ip1a
supramolecular fiber
katnal2
kif5aa
ttll6
dusp27
kif26bb
si:ch211−114n24.6
dnal1
hmcn2
unc45b
myom1a
hcls1
mdm1
myoz1b
col5a3a
col11a1b
extracellular matrix
spon2a
lamc3
spon1b
lama5
ntn2
tecta
ntn1a
bcan
ccn1l2
emilin2a
lamb1a
lama1
lama4
col7a1
fgf3
col8a1a
col28a1a
matn1
col17a1b
si:dkeyp−44a8.4
mmp30
mmp14a
bmper
zgc:123295
c3b.1
rspo3
bmp8a
loxl3a
smpdl3b
lft2
lft1
prss59.1
dkk1b
si:ch1073−459j12.1
sema3bl
ndr1
ifnphi1
apoa4b.2
extracellular space
serpini1
ces2b
pcsk1
sfrp2
ctsl.1
anxa1b
wnt11f2
fbln5
anxa1c
tgfb5
fgf8a
metrn
cd109
bmp7b
serpina10a
frzb
tgfb1a
angptl4
ndr2
serpinb1l4
pyyb
tnfb
fetub
wnt5b
apoc2
zgc:163030
sema4e
ifnphi3
f10
gprc5bb
masp1
tnfaip6
sostdc1a
size
13
29
44
60
fold change
12
9
6
3

### Figure S2

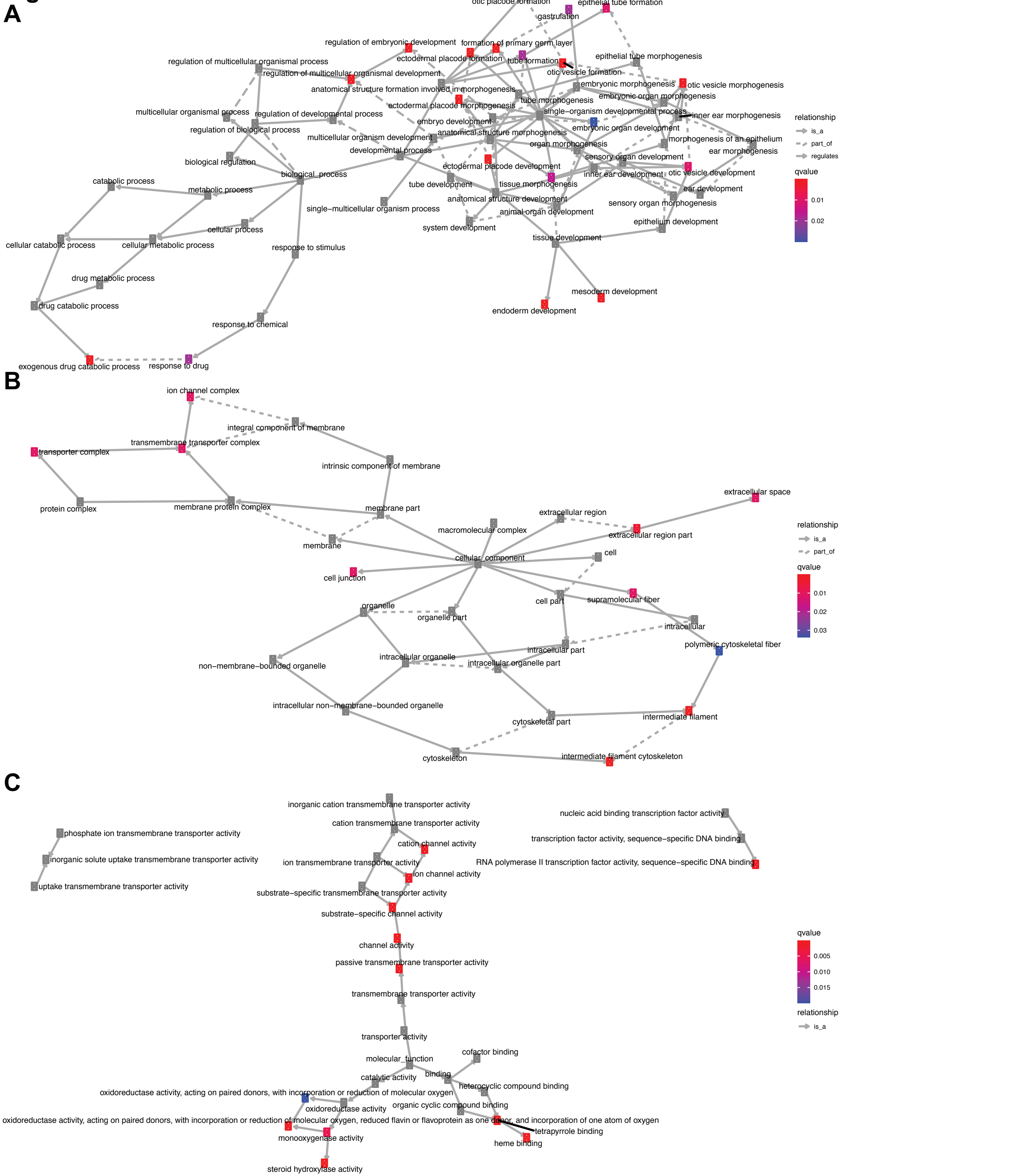
A
otic placode formation
epithelial tube formation
gastrulation
regulation of embryonic development
formation of primary germ layer
ectodermal placode formation
tube formation
epithelial tube morphogenesis
regulation of multicellular organismal process
regulation of multicellular organismal development
otic vesicle formation
embryonic morphogenesis
otic vesicle morphogenesis
anatomical structure formation involved in morphogenesis
embryonic organ morphogenesis
tube morphogenesis
ectodermal placode morphogenesis
single−organism developmental process
multicellular organismal process
regulation of developmental process
inner ear morphogenesis
embryo development
embryonic organ development
regulation of biological process
anatomical structure morphogenesis
multicellular organism development
morphogenesis of an epithelium
organ morphogenesis
developmental process
ear morphogenesis
sensory organ development
biological regulation
ectodermal placode development
inner ear development
otic vesicle development
biological_process
catabolic process
tube development
tissue morphogenesis
metabolic process
ear development
anatomical structure development
single−multicellular organism process
sensory organ morphogenesis
animal organ development
cellular process
system development
epithelium development
cellular catabolic process
cellular metabolic process
tissue development
response to stimulus
drug metabolic process
mesoderm development
drug catabolic process
endoderm development
response to chemical
exogenous drug catabolic process
response to drug
relationship
is_a
part_of
regulates
qvalue
0.01
0.02
B
ion channel complex
integral component of membrane
transmembrane transporter complex
transporter complex
intrinsic component of membrane
protein complex
membrane protein complex
membrane part
extracellular space
extracellular region
macromolecular complex
extracellular region part
membrane
cellular_component
cell
cell junction
organelle
cell part
supramolecular fiber
organelle part
intracellular
intracellular part
polymeric cytoskeletal fiber
non−membrane−bounded organelle
intracellular organelle
intracellular organelle part
intracellular non−membrane−bounded organelle
cytoskeletal part
intermediate filament
cytoskeleton
intermediate filament cytoskeleton
relationship
is_a
part_of
qvalue
0.01
0.02
0.03
C
inorganic cation transmembrane transporter activity
nucleic acid binding transcription factor activity
phosphate ion transmembrane transporter activity
cation transmembrane transporter activity
transcription factor activity, sequence−specific DNA binding
cation channel activity
inorganic solute uptake transmembrane transporter activity
ion transmembrane transporter activity
RNA polymerase II transcription factor activity, sequence−specific DNA binding
ion channel activity
uptake transmembrane transporter activity
substrate−specific transmembrane transporter activity
substrate−specific channel activity
channel activity
passive transmembrane transporter activity
transmembrane transporter activity
transporter activity
molecular_function
cofactor binding
catalytic activity
binding
heterocyclic compound binding
oxidoreductase activity, acting on paired donors, with incorporation or reduction of molecular oxygen
oxidoreductase activity
organic cyclic compound binding
oxidoreductase activity, acting on paired donors, with incorporation or reduction of molecular oxygen, reduced flavin or flavoprotein as one donor, and incorporation of one atom of oxygen
tetrapyrrole binding
monooxygenase activity
heme binding
steroid hydroxylase activity
qvalue
0.005
0.010
0.015
relationship
is_a

### Figure S3

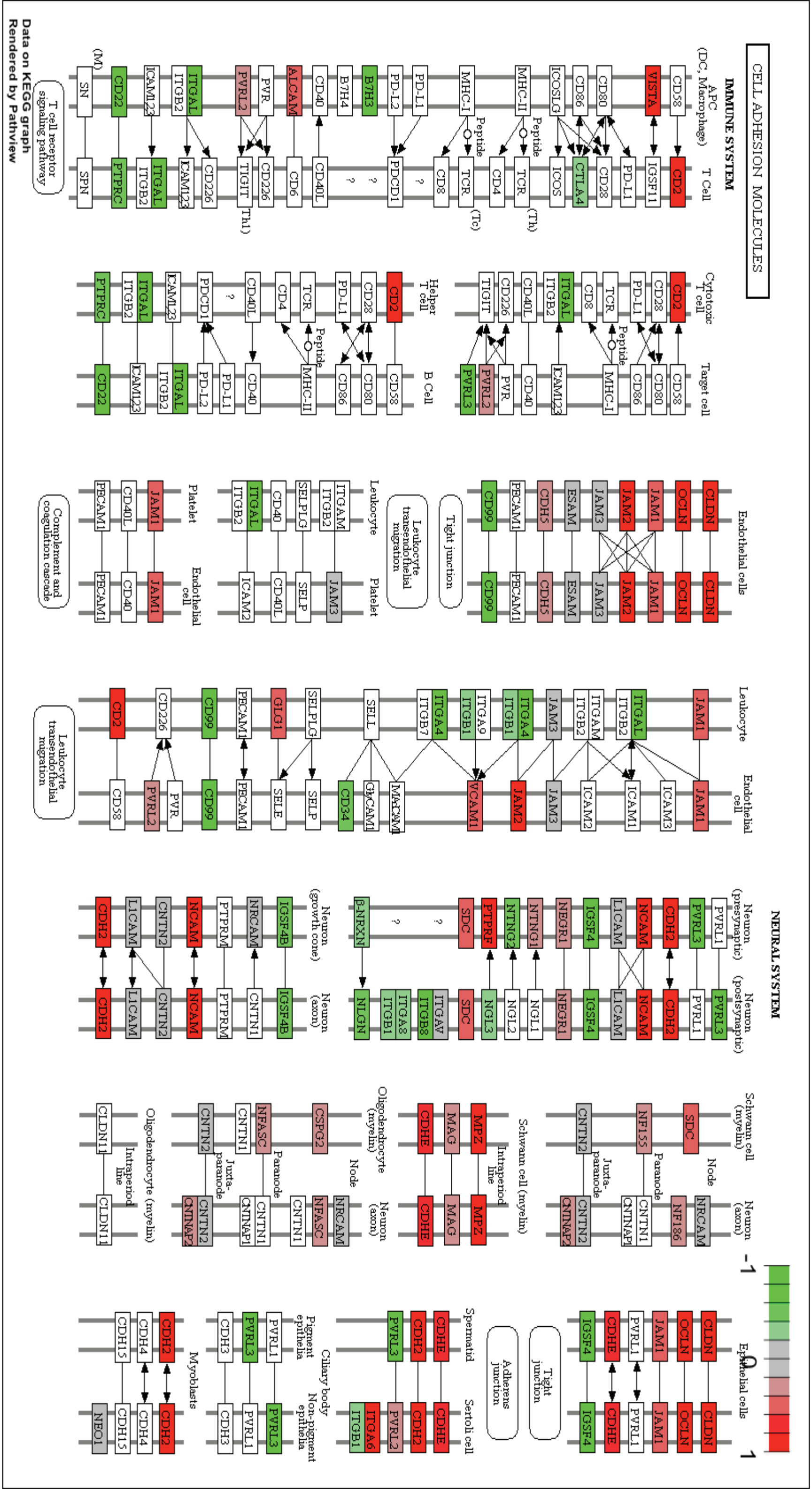

CELL ADHESION MOLECULES
IMMUNE SYSTEM
NEURAL SYSTEM
APC (DC, Macrophage)
T Cell
Cytotoxic T cell
Target cell
Helper T cell
B Cell
Endothelial cells
Leukocyte
Platelet
Endothelial cell
Tight junction
Leukocyte transendothelial migration
Complement and coagulation cascade
T cell receptor signaling pathway
Neuron (presynaptic)
Neuron (postsynaptic)
Neuron (growth cone)
Neuron (axon)
Schwann cell (myelin)
Oligodendrocyte (myelin)
Node
Paranode
Juxta-paranode
Intraperiod line
Epithelial cells
Adherens junction
Spermatid
Sertoli cell
Ciliary body
Pigment epithelia
Non-pigment epithelia
Myoblasts
Peptide
VISTA
CD58
CD80
CD86
ICOSLG
MHC-II
MHC-I
PD-L1
PD-L2
B7H3
B7H4
CD40
ALCAM
PVR
PVRL2
ITGAL
ITGB2
ICAM123
CD22
SN
CD2
IGSF11
CD28
CTLA4
ICOS
TCR
CD4
CD8
PDCD1
CD40L
CD6
CD226
TIGIT
PTPRC
SPN
CLDN
OCLN
JAM1
JAM2
JAM3
ESAM
CDH5
PECAM1
CD99
ITGAM
SELPLG
SELP
ICAM2
ICAM1
ICAM3
ITGA4
ITGB1
ITGA9
ITGB7
SELL
GLG1
SELE
CD34
GlyCAM1
MAdCAM1
VCAM1
PVRL1
PVRL3
CDH2
NCAM
L1CAM
IGSF4
NEGR1
NTNG1
NTNG2
PTPRF
SDC
NGL1
NGL2
NGL3
ITGAV
ITGB8
ITGA8
NLGN
β-NRXN
IGSF4B
NRCAM
PTPRM
CNTN1
CNTN2
NF155
NF186
CNTNAP1
CNTNAP2
MPZ
MAG
CDHE
CSPG2
NFASC
CLDN11
ITGA6
CDH3
CDH4
CDH15
NEO1
-1
0
1
Data on KEGG graph
Rendered by Pathview

### Figure S4

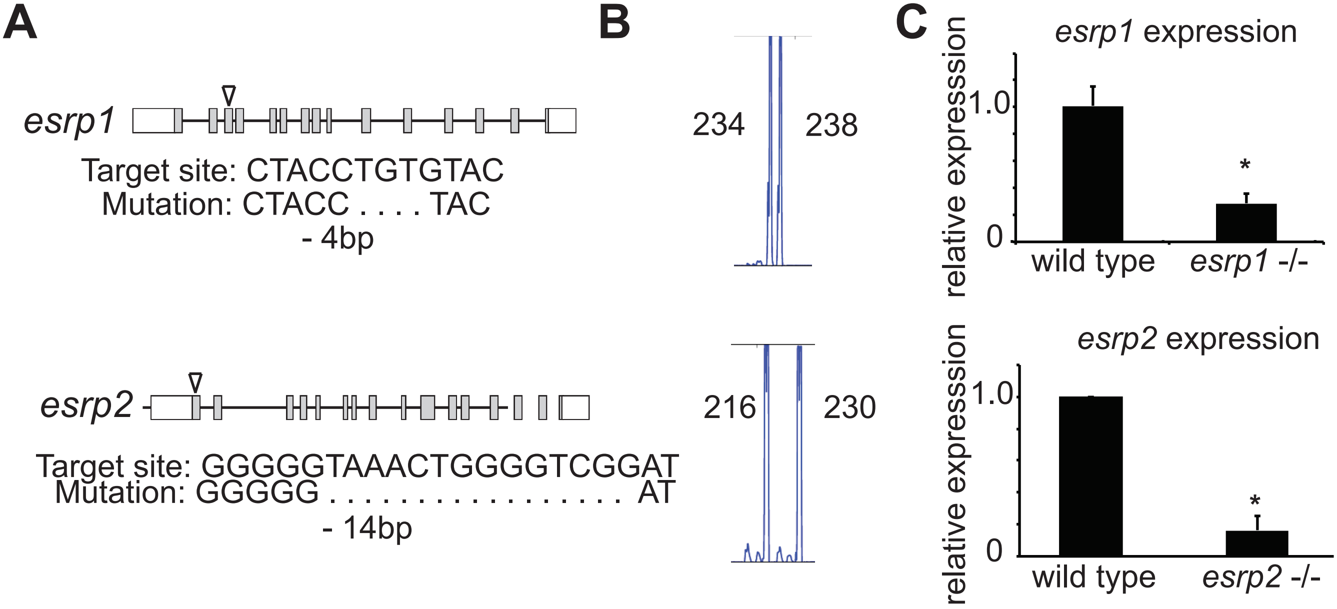
A
esrp1
Target site: CTACCTGTGTAC
Mutation: CTACC . . . . TAC
- 4bp
esrp2
Target site: GGGGGTAAACTGGGGTCGGAT
Mutation: GGGGG . . . . . . . . . . . . . . . . . . AT
- 14bp
B
234
238
216
230
C
esrp1 expression
relative expression
1.0
0
*
wild type
esrp1 -/-
esrp2 expression
relative expression
1.0
0
*
wild type
esrp2 -/-

| Table S1. Irf6, Esrp1 and Esrp2 genotypes interact to produce non-Mendelian embryo ratios | | | | | |
|---|---|---|---|---|---|
| ***Irf6*** | ***Esrp1*** | ***Esrp2*** | **probability** | **Expected** | **Observed** |
| WT | WT | WT | 1/64 | 0.8 | 1 |
| WT | WT | Het | 1/32 | 1.5 | 2 |
| WT | WT | KO | 1/64 | 0.8 | 1 |
| WT | Het | WT | 1/32 | 1.5 | 2 |
| WT | Het | Het | 1/16 | 3.1 | 2 |
| WT | Het | KO | 1/32 | 1.5 | 3 |
| WT | KO | WT | 1/64 | 0.8 | 0 |
| WT | KO | Het | 1/32 | 1.5 | 1 |
| WT | KO | KO | 1/64 | 0.8 | 1 |
| Het | WT | WT | 1/32 | 1.5 | 3 |
| Het | WT | Het | 1/16 | 3.1 | 4 |
| Het | WT | KO | 1/32 | 1.5 | 1 |
| **Het** | **Het** | **WT** | 1/16 | 3.1 | 7 |
| Het | Het | Het | 1/8 | 6.1 | 3 |
| **Het** | **Het** | **KO** | 1/16 | 3.1 | 0 |
| Het | KO | WT | 1/32 | 1.5 | 1 |
| Het | KO | Het | 1/16 | 3.1 | 5 |
| Het | KO | KO | 1/32 | 1.5 | 0 |
| KO | WT | WT | 1/64 | 0.8 | 0 |
| KO | WT | Het | 1/32 | 1.5 | 2 |
| KO | WT | KO | 1/64 | 0.8 | 0 |
| KO | Het | WT | 1/32 | 1.5 | 3 |
| KO | Het | Het | 1/16 | 3.1 | 3 |
| KO | Het | KO | 1/32 | 1.5 | 3 |
| KO | KO | WT | 1/64 | 0.8 | 0 |
| KO | KO | Het | 1/32 | 1.5 | 0 |
| KO | KO | KO | 1/64 | 0.8 | 0 |

*Irf6R84C/WT*; *Esrp1*+/-; *Esrp2*+/- triple heterozygous mice were in-crossed and embryos were collected between E12.5 and E21. Table 1 is a subset of expected number of embryos based on Mendelian genetics versus the observed number of viable embryos. The *Irf6R84C/WT*; *Esrp1*+/-; *Esrp2*-/- genotype appears to be lethal prior to E12.5 as approximately 3 embryos were expected but zero embryos were observed.  A total of 49 embryos were collected from 7 different litters.
